# Structural Basis of Lipopolysaccharide Assembly by the Outer Membrane Translocon Holo-Complex

**DOI:** 10.1101/2025.04.10.648201

**Authors:** Haoxiang Chen, Axel Siroy, Violette Morales, Dominik Gurvic, Yves Quentin, Stephanie Balor, Yassin A. Abuta’a, Maurine Marteau, Carine Froment, Anne Caumont-Sarcos, Julien Marcoux, Phillip J. Stansfeld, Rémi Fronzes, Raffaele Ieva

## Abstract

Lipopolysaccharide (LPS) assembly at the surfaces-exposed leaflet of the bacterial outer membrane (OM) is mediated by the OM LPS translocon. An essential transmembrane β-barrel protein, LptD, and a cognate lipoprotein, LptE, translocate LPS selectively into the OM external leaflet via a poorly understood mechanism. Here, we characterize two additional translocon subunits, the lipoproteins LptM and LptY (formerly YedD). We use single-particle cryo-EM analysis, functional assays and molecular dynamics simulations to visualize the roles of LptM and LptY at the translocon holo-complex LptDEMY, uncovering their impact on LptD conformational dynamics. Whereas LptY binds and stabilizes the periplasmic LptD β-taco domain that functions as LPS receptor, LptM intercalates the lateral gate of the β-barrel domain, promoting its opening and access by LPS. Remarkably, we demonstrate a conformational switch of the LptD β-taco/β-barrel interface alternating between contracted and extended states. The LptD β-strand 1, which defines the mobile side of the lateral gate, binds LPS and performs a stroke movement toward the external leaflet during the contracted-to-extended state transition. Our findings establish a detailed mechanistic framework explaining the selective transport of LPS to the membrane external leaflet.

## INTRODUCTION

Diderm bacteria are encased by an outer membrane (OM) that serves as a protective barrier and a platform for environmental interactions ^1,2^. The OM is typically characterized by the presence of integral outer membrane proteins (OMPs), which fold into transmembrane β-barrels ^3^. Some OMPs play crucial roles in OM biogenesis and molecular transport. Among these, the OMP foldase BamA (also known as subunit A of the β-barrel assembly machinery, or BAM) and the transporter LptD are hallmark OMPs found in the vast majority of diderm bacteria, including both those traditionally classified as diderm Gram-negative (*e.g.* Proteobacteria) and diderm Gram-positive (*e.g.* Negativicutes) ^4–6^. In Proteobacteria, LptD, together with its cognate lipoprotein LptE, forms the OM translocon responsible for inserting lipopolysaccharide (LPS) into the external leaflet ^7–10^. This process establishes an asymmetric bilayer, with glycerophospholipids confined to the internal leaflet ^11–13^. The asymmetric distribution of LPS is crucial for organizational and structural roles of the OM, making it essential for the survival of most Gram-negative bacteria. Neighbouring LPS molecules interact electrostatically via phosphate groups that are bridged by divalent cations. This tightly packed molecular arrangement forms a membrane structure that is not only impermeable to many antibiotics and surfactants, but also resistant to physical stress ^14,15^.

LPS has a tripartite structure, made of lipid A, the core oligosaccharide, and the O-antigen. Lipid A is the hydrophobic moiety, comprising 4 to 7 acyl tails attached to a disaccharide of two phosphorylated N-acetyl glucosamine units. The headgroup of lipid A is decorated with a core oligosaccharide of approximately 10 sugars, which can be further extended with the addition of the O-antigen, a highly variable sugar segment composed of repeated oligosaccharide units ^16–18^. The most conserved portion of LPS is Kdo-lipid A (KLA) consisting of lipid A and two 3-deoxy-D-manno-oct-2-ulosonic acid (Kdo) sugars from the oligosaccharide core ^19^. The remaining saccharide components are generally dispensable in *Escherichia coli* under laboratory growth conditions. Anterograde LPS transport (Lpt) from its site of biosynthesis at the inner membrane (IM) to the OM is mediated by the 7-component Lpt pathway, which forms a transenvelope bridge ^7,20^. An ATP-binding cassette (ABC) transporter, LptB_2_FG, extracts LPS from the IM and transfers it to the IM-anchored protein LptC. LptA then ferries LPS across the periplasm, from LptC to the OM translocon LptDE. During transport, the lipid tails of LPS move along the hydrophobic grooves formed by stacked β-taco domains ^21^, including those of LptFG and LptC at the IM, LptA in the periplasm, and LptD at the OM ^22,23^ (Fig. 1a).

**Figure 1.**
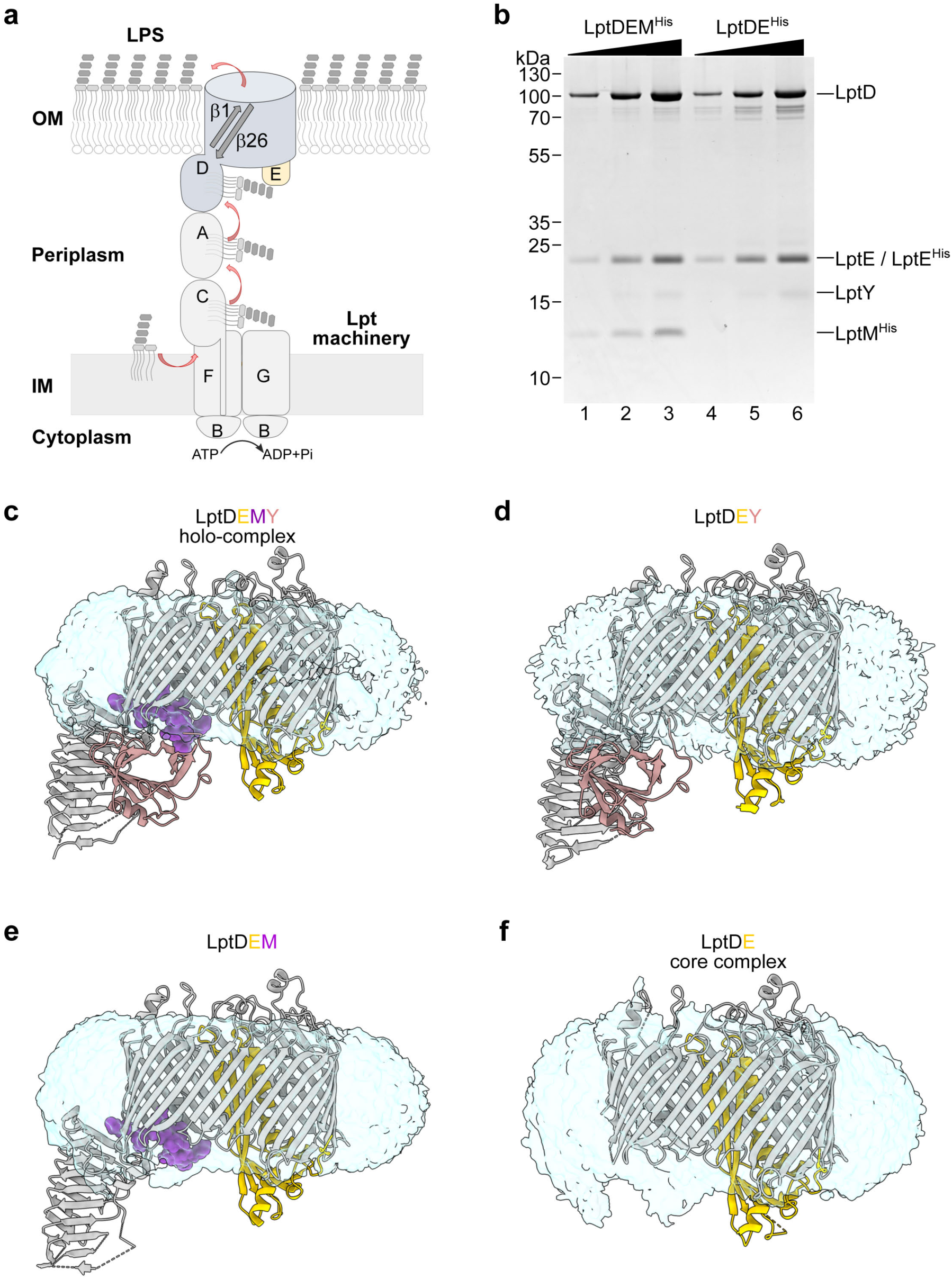
Purification and structural determination of OM LPS translocon complexes. **a**, Schematic representation of the Lpt pathway. **b**, Incremental amounts of the indicated purified translocons were loaded on SDS-PAGE and revealed by Coomassie Blue staining. **c-f**, Cryo-EM structures of the LptDEMY translocon holo-complex or the indicated subcomplexes in DDM micelles. LptD (grey), LptE (yellow), and LptY (pink) are shown in ribbon representation, whereas LptM (purple) is shown in sphere representation.

The transmembrane domain of LptD forms a β-barrel composed of 26 β-strands (designated β1 to β26), with an internal lumen partially occluded by the lipoprotein LptE ^24–27^. The β-barrel domain of LptD is folded around LptE at the BAM complex ^28^. Covalent bonds connect the β-taco domain to both the N- and C-terminal portions of the β-barrel domain ^29–31^. The polypeptide backbone linking the β-taco to β1 and the double-disulfide bond anchoring the β-taco to the periplasmic turn β24-β25 create a conduit from the periplasm to the OM. It has been proposed that the acyl tails of LPS enter the membrane near the LptD β-taco/β-barrel interface region, whereas the LPS polysaccharides transverse the β-barrel lumen adjacent to LptE and exits at the cell surface through a putative lateral gate between the terminal β-barrel strands, β1 and β26 ^24,25,32,33^ (Fig. 1a). Experimental evidence supports possible opening of the LptD lateral gate in agreement with this model ^25,34,35^. Nevertheless, all high-resolution structures have captured the LptD lateral gate in a closed conformation. Complete separation of the lateral gate strands has been observed only upon molecular dynamics (MD) simulations under conditions of high negative pressure ^25^ or upon binding with macrobodies ^35^. Consequently, the precise mechanism by which LPS is selectively released into the OM external leaflet remains unclear.

We recently identified the lipoprotein LptM as a new stoichiometric component of the OM LPS translocon ^36^. Our findings demonstrated that, like LptE, LptM associates with LptD during the folding of its transmembrane domain at the BAM complex, a step that facilitates subsequent disulfide bond formation between the β-barrel and β-taco domains ^36^. However, whether LptM plays a role in LPS translocation has remained unknown. In this study, while determining the LptDEM structure, we characterize an additional component of the translocon, the lipoprotein YedD that we have renamed LptY based on the Lpt nomenclature ^37^. Most importantly, functional assays and structural analyses of the translocon holo-complex LptDEMY and of its subcomplexes, LptDEM, LptDEY and LptDE, reveal that LptD operates through a conformational switch between contracted and extended states of its β-taco/β-barrel interface near the β1-β26 lateral gate. Whereas LptY stabilizes the LptD β-taco domain, LptM is positioned at the lateral gate facilitating its opening and access by LPS.

## RESULTS

### The OM LPS translocon interacts with LptY (formerly YedD)

To investigate whether LptM plays a role in LPS translocation, we aimed at determining the structures of LptDE translocon with and without LptM. We purified both LptDEM and the core translocon LptDE via nickel-affinity chromatography and gel filtration. As baits, we used respectively LptM^His^ expressed in wild-type cells or LptE^His^ expressed in Δ*lptM* cells (Fig. 1b). Notably, both eluates contained an additional protein migrating at an apparent molecular mass of approximately 15 kDa. Bottom-up mass spectrometry on the corresponding gel bands identified this protein as LptY, an OM lipoprotein of ∼14 kDa ^38^ (Extended Data Fig. 1a, b). Phylogenetic analysis showed that, like *lptM*, *lptY* is conserved within *Enterobacteriaceae* (Extended Data Fig. 1c) and *Erwiniaceae,* except for species presenting highly reduced genomes, which often lack also *lptD and lptE*. Furthermore, *lptY* is present in some *Pectobacteriaceae,* but is absent in other families of *Enterobacterales* such as *Yersiniaceae*, *Hafniaceae*, *Morganellaceae* and *Budviciaceae*, which retain *lptM* along with *lptD* and *lptE* (Extended Data Figs. 2 and 3, see also Supplementary Methods).

In our previous study, native mass-spectrometry analysis of LptDEM and LptDE revealed respectively 1:1:1 and 1:1 stoichiometries ^36^. Revisiting these spectra in consideration of LptY’s identification, we detected a subpopulation of both LptDEM and LptDE with a mass shift of 14,301 Da, consistent with complexes bound to LptY. This yielded LptY-bound complexes with 1:1:1:1 and a 1:1:1 stoichiometries (Extended Data Fig. 4; see also in Ref ^36^ and the PRIDE repository dataset PXD041774). MS spectra obtained in the low m/z region identified a species at 14,302 Da, corresponding to LptY dissociated from LptDEM^His^ and LptDE^His^ complexes (Extended Data Fig. 4b and d). Supporting these findings, LptY pull-down experiments co-isolated LptD and LptM, indicating stable association with the LPS translocon (Extended Data Fig. 5). Thus, LptY emerges as a new component of the OM LPS translocon capable of binding LptDE alongside LptM. We refer to LptDE bound to both LptM and LptY as the LPS translocon “holo-complex”.

### Structural determination of the LPS translocon holo-complex and sub-complexes

Using cryo-EM and single particle analysis, we determined the structures of the purified OM LPS translocon complexes in n-dodecyl-β-D-maltopyranoside (DDM) micelles (Fig. 1c-f and Extended Data Figs. 6 and 7). The structure of the core complex, LptDE, was solved at an overall resolution of 2.74 Å for the transmembrane portion including the LptD β-barrel and LptE. However, in the reconstructed maps, no density was visible for the LptD β-taco domain (Fig. 1f and Extended Data Fig. 7d), indicating conformational flexibility of this domain as previously documented ^24,34^. Flexibility of the β-taco domain in the absence of LptM was also suggested by our previous hydrogen-deuterium exchange (HDX)-MS results, showing high deuteration levels for the β-taco in the LptDE core complex ^36^. Furthermore, we solved the structures of LptDE bound to either LptM, LptY, or both proteins (holo-complex) at resolutions of 2.47, 2.62 and 2.63 Å, respectively. In these complexes, binding of LptM, LptY, or both stabilized the β-taco domain, enabling its structural determination (Fig. 1c-e and Extended Data Fig. 7a-c).

### LptM intercalates the LptD β-taco/β-barrel lateral gate interface

The electron density map allowed *de novo* model building of the N-terminal portion of LptM, including the first 10 amino acid residues of the mature sequence following its N-terminal Cys residue (LptM_20-29_) (Fig. 2a). This segment is part of the Pfam motif PF13627, which is highly conserved across bacterial species harbouring *lptM* ^36^. The LptM N-terminal region alone (LptM^1144–67^) is sufficient to mediate its co-purification with the translocon (Extended Data Fig. 8a). Conversely, an internal truncation of residues 22-30 of LptM (which includes a significant portion of the N-terminal part resolved in our cryo-EM structure, LptM^1122–30^) strongly impairs association with the translocon (Extended Data Fig. 8a). The structurally resolved portion of LptM forms extensive contacts with the internal surface of the LptD β-barrel, particularly with residues located at the periplasmic-facing side of β1-β4, as well as with the β-taco/β-barrel hinge region preceding β1 (Fig. 2b). This interaction between LptD and LptM remains largely unchanged in the absence of LptY (Extended Data Fig. 8b and c; Extended Data Table 1). In both LptDEM and LptDEMY structures, we resolved portions of the LptM lipid tails in positions proximal to the external surface of the LptD lateral gate. Notably, one of the acyl tails intercalates between the outward-facing side chains of β1 and β26 (Fig. 2a and Extended Data Fig. 8b).

**Figure 2.**
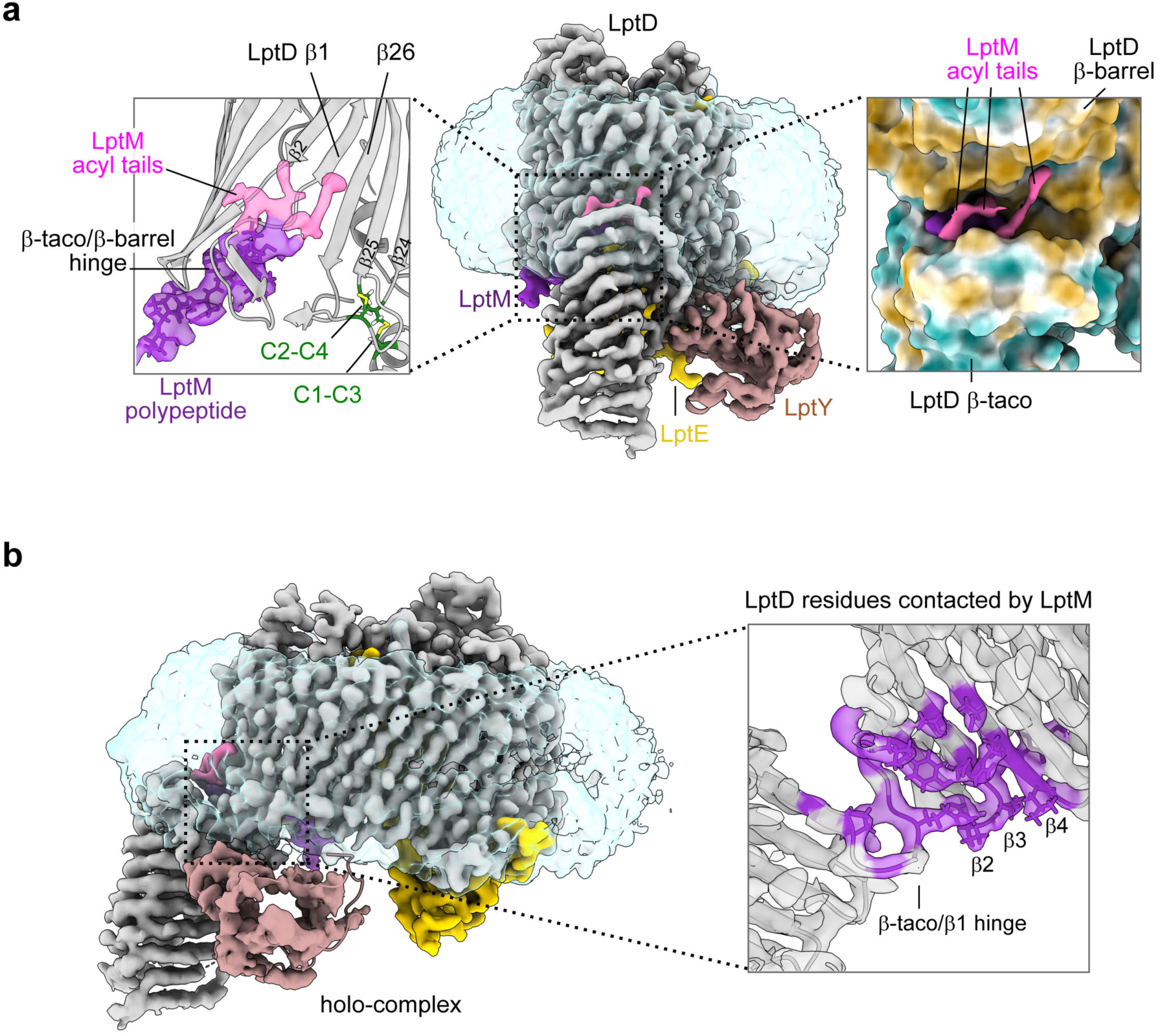
Interaction of LptM with the LptD β-barrel/β-taco interface. **a**, **Left**, Zoom on LptM in the structure of LptDEMY, highlighting the electron density corresponding to the acyl tails of LptM (pink) and the electron density corresponding to the LptM N-terminal moiety (LptM_20-29_, purple) situated near the lateral gate of LptD (β1-β26). The β-taco/β-barrel hinge and the LptD inter-domain disulfide bonds surrounding LptM are indicated. LptD Cys31 (C1), Cys173 (C2), Cys724 (C3) and Cys725 (C4) are shown in green with disulfide bonds in yellow. **Central**, cryo-EM map of the LptDEMY complex coloured according to the protein chains (LptD in grey, LptE in yellow, LptM in purple, LptY in pink). **Right**, Zoom on the acyl tails of LptM and the surrounding surface of LptD coloured based on hydrophobicity of the amino acid side chains (brown, hydrophobic; cyan, hydrophilic). **b**, Internal section of the LptD β-taco/β-barrel hinge region in the holo-complex. The LptD residues in contact with LptM are shown in purple.

The LptD β-taco domain is anchored to both sides of the β-barrel lateral gate. On the N-terminal side, the polypeptide backbone connects the β-taco to β1. On the C-terminal side Cys31 (C1) and Cys173 (C2) in the β-taco are linked via disulfide bonds respectively to Cys724 (C3) and Cys725 (C4) located in the β24-β25 periplasmic turn (Fig. 2a) ^29^. Positioned at the β1-β26 lateral gate, LptM is constrained between the two interdomain connections and the inner leaflet of the membrane, thus raising the question of how LptM reaches its location in the translocon. We had previously shown that LptM is assembled with LptD at the BAM complex ^36^, where an early oxidation intermediate of LptD lacking interdomain disulfides undergoes folding ^30^. It is therefore likely that the N-terminus of LptM is positioned within the translocon before the β-taco and the β-barrel domains are connected by disulfide bonds ^36^.

### LptY interacts with LptD β-taco domain independently of LptM

The cryo-EM density map revealed clear features corresponding to LptY adjacent to the LptD β-taco domain. We generated a predicted structure of *E. coli* LptY using AlphaFold 2 ^39^, which were consistent with the previously determined X-ray crystal structure of the non-lipidated variant of its ortholog from *Klebsiella pneumoniae* (PDB: 4HWM). The predicted AlphaFold 2 model could be fitted into the experimental density using rigid body docking, despite its limited resolution in this region of the map (Fig. 3a and Extended Data Fig. 9a). LptY adopts an 8-stranded β-barrel fold flanked by a short N-terminal 3_10_-helix and a C-terminal α-helix. This organization of structural motifs resembles that of lipocalins, a family of proteins capable of interacting with hydrophobic ligands ^40^. Notably, LptY contains two conserved cysteine residues (Extended Data Fig. 1c), one in β-strand 3 and one in the C-terminal α-helix), which are positioned at a distance compatible with disulfide bond formation (Extended Data Fig. 9a). Structural analysis of the two LptY-containing complexes revealed similar contact areas with the external surface of LptD β-taco domain and its N-terminal α-helix (Fig. 3a and Extended Data Table 2). During our study, another paper has described the cryo-EM structure of the *E. coli* LPS translocon bound to LptY, showing a similar interaction between the β-taco domain of LptD and LptY ^41^. Indeed, our pull-down of LptY^His^ from cells co-expressing endogenous LptD and an N-terminally truncated version lacking the β-taco domain, LptD^β-barrel^, only co-eluted full-length LptD, thus proving the critical role of the β-taco domain for LptY binding. In contrast, pull-down of LptM^His^, which interacts with the LptD β-barrel domain, co-eluted both the full-length and the N-terminally truncated β-barrel forms of LptD (Extended Data Fig. 9b) ^36^. The independent binding of LptD to LptM and LptY is further highlighted by the fact that we could determine the structures of the core translocon LptDE bound to either one of the newly identified translocon lipoproteins (Fig. 1d and e).

**Figure 3.**
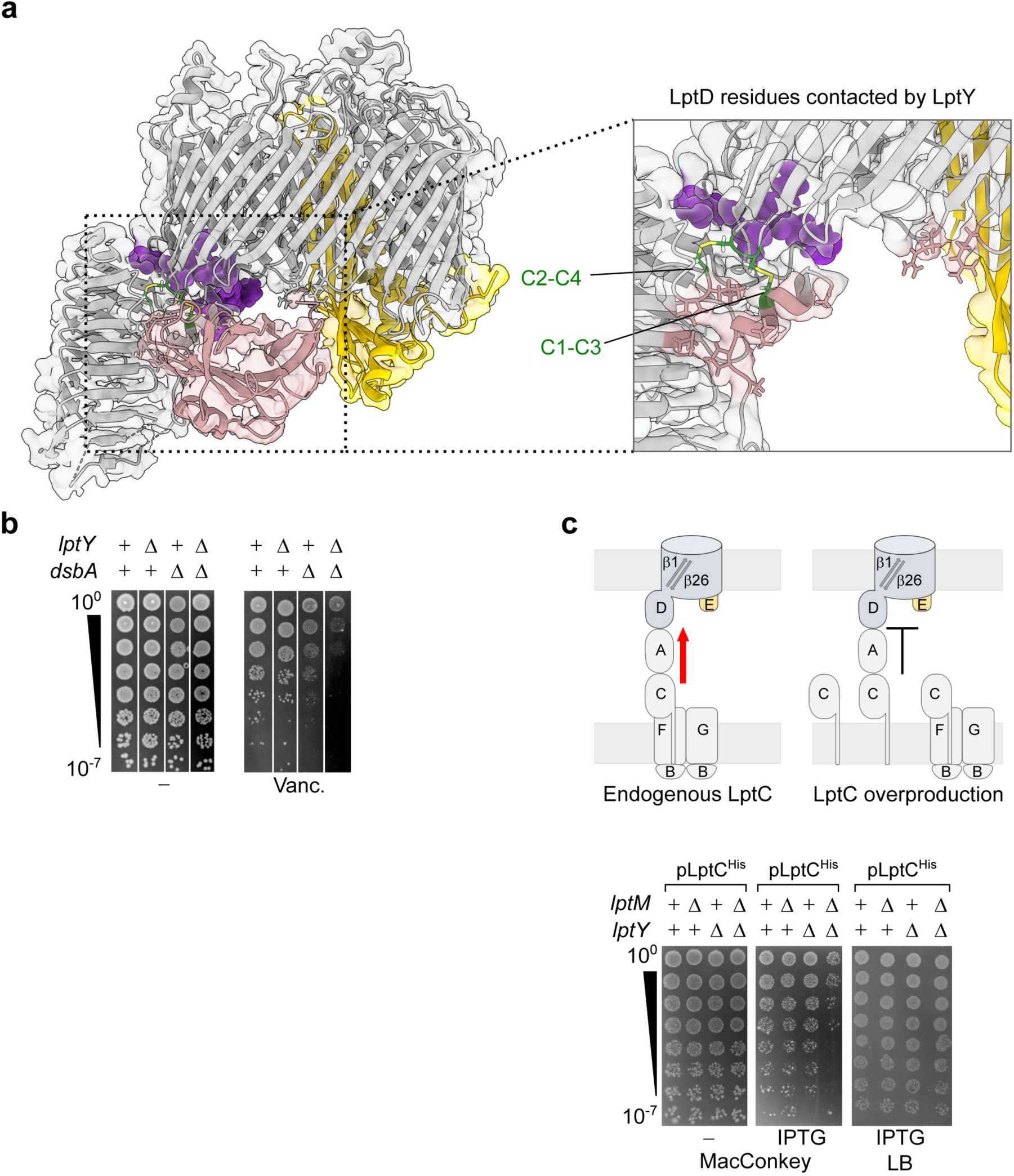
Interaction of LptY with the LptD β-taco domain. **a**, Zoom on the LptD area in contact with LptY, proximal to the LptD inter-domain disulfide bonds. Proteins shown in ribbon representation are LptD (grey), LptE (yellow), and LptY (pink). LptM is shown in sphere representation (purple). **b**, Drop dilution test of the indicated strains on regular LB agar media or LB agar supplemented with vancomycin. **c, Top**, Schematics of the predicted effects of LptC overproduction on Lpt bridges. **Bottom**, drop dilution spot test of the indicated strains on MacConkey agar media or on regular LB agar media, as indicated.

Given LptY’s interaction proximal to a region of LptD that undergoes oxidative maturation, we investigated whether LptY plays a role in LptD biogenesis. Deletion of *lptY* does not affect the levels of oxidized LptD (Extended Data Fig. 9c), nor leads to accumulation of LptDE at the BAM complex (Extended Data Fig. 9d). Interestingly, we found LptY to be critical for OM integrity against the large-molecular weight antibiotic vancomycin in cells lacking the cysteine oxidase *dsbA* (Fig. 3b). DsbA catalyzes LptD oxidation ^29^, and cells lacking this enzyme show reduced LptD levels associated with mild sensitivity to vancomycin ^36^. Conversely, Δ*lptY* cells were not sensitive to the tested antibiotic concentrations. However, deleting *lptY* in Δ*dsbA* resulted in a synergistic detrimental effect that further enhanced OM permeability compared to Δ*dsbA* alone (Fig. 3b), highlighting the importance of LptY when LptD levels are reduced.

Similar to LptM, which we showed to associate with IM and periplasmic Lpt components ^36^, LptY purification co-eluted periplasmic LptA (Extended Data Fig. 5), indicating that both newly identified components are part of transenvelope Lpt bridges. Because our structures showed that LptM and LptY stabilize the LptD β-taco domain, which connects the OM and IM portions of the Lpt bridge, we investigated whether cells lacking these newly identified components would be more sensitive to perturbations of Lpt bridges. To explore this hypothesis, we examined the effects of overproducing LptC. This condition was shown to titrate the OM translocon via LptA by forming incomplete bridges because of the excess LptC relative to LptB_2_FG ^22^. We reasoned that reducing the number of complete Lpt bridges would make cells more dependent on efficient functioning of the remaining IM-OM Lpt connections (Fig. 3c, diagram). Upon LptC overproduction, cell susceptibility to bile salts (MacConkey media) increased, especially in a strain harbouring both deletions of *lptM* and *LptY* (Fig. 3c). This result suggests that the newly identified OM LPS translocon components contribute to optimal functioning of Lpt bridges.

### LptD functions through the contraction and extension of its β-taco/β-barrel interface

Our structural analysis has revealed that both LptM and LptY contribute to stabilizing the LptD β-taco domain, enabling its structural determination. To explore other potential variations within the translocon holo-complex, its variants lacking either LptY or LptM, and the core complex, we superimposed the four complexes and assessed the root mean square deviation (RMSD) of atomic distances. The membrane portion of the core translocon was superimposable with that of the holo-complex, resulting in an RMSD lower than 0.1 Å for the largest portion of LptE and the LptD β-barrel domain (Extended Data Fig. 10a and d). Similarly, the holo-complex and the LptDEM complex showed an RMSD lower than 0.1 Å for the membrane-embedded portion of the translocon and an RMSD ranging from 0.1 to 0.8 Å for the β-taco domain (Extended Data Fig. 10a and b).

Remarkably, superimposition of the holo-complex with the variant lacking LptM (LptDEY) revealed significant deviations in the β-barrel lobe containing the lateral gate, with > 2 Å deviations for β1-β4 and β26 (Extended Data Fig. 10a and c). These conformational variations in the LptD β-barrel domain result from a +2 increment in the β-barrel shear number ^42^. This means that β1 slides along β26 of two amino acid positions towards the periplasm (Fig. 4a). Simultaneously, the hinge loop at the N-terminal side of β1 bends toward the β-barrel lumen, causing the β-taco to move toward the membrane (Fig. 4a and Extended Data Fig. 11a). The convergence of the β-taco and β1 movement reduces the gap between the two LptD domains (measured between residues P219 in the β-taco and N232, the first residue of β1) from 19.7 Å in presence of LptM to 12.0 Å in its absence (Fig. 4b). Henceforth we will name these distinct LptD conformations “contracted” state, as observed in the structure of LptDEY, and “extended” state, as observed in LptDEMY and LptDEM structures. The structural deviations observed at the β-taco/β-barrel interface in the absence of LptM fit well with the bimodal distribution of hydrogen-deuterium exchange (HDX) kinetics that we have previously documented using HDX-mass spectrometry on the same sample. LptM, instead, abrogates this bimodal behaviour, stabilizing the interface between the two domains ^36^.

**Figure 4.**
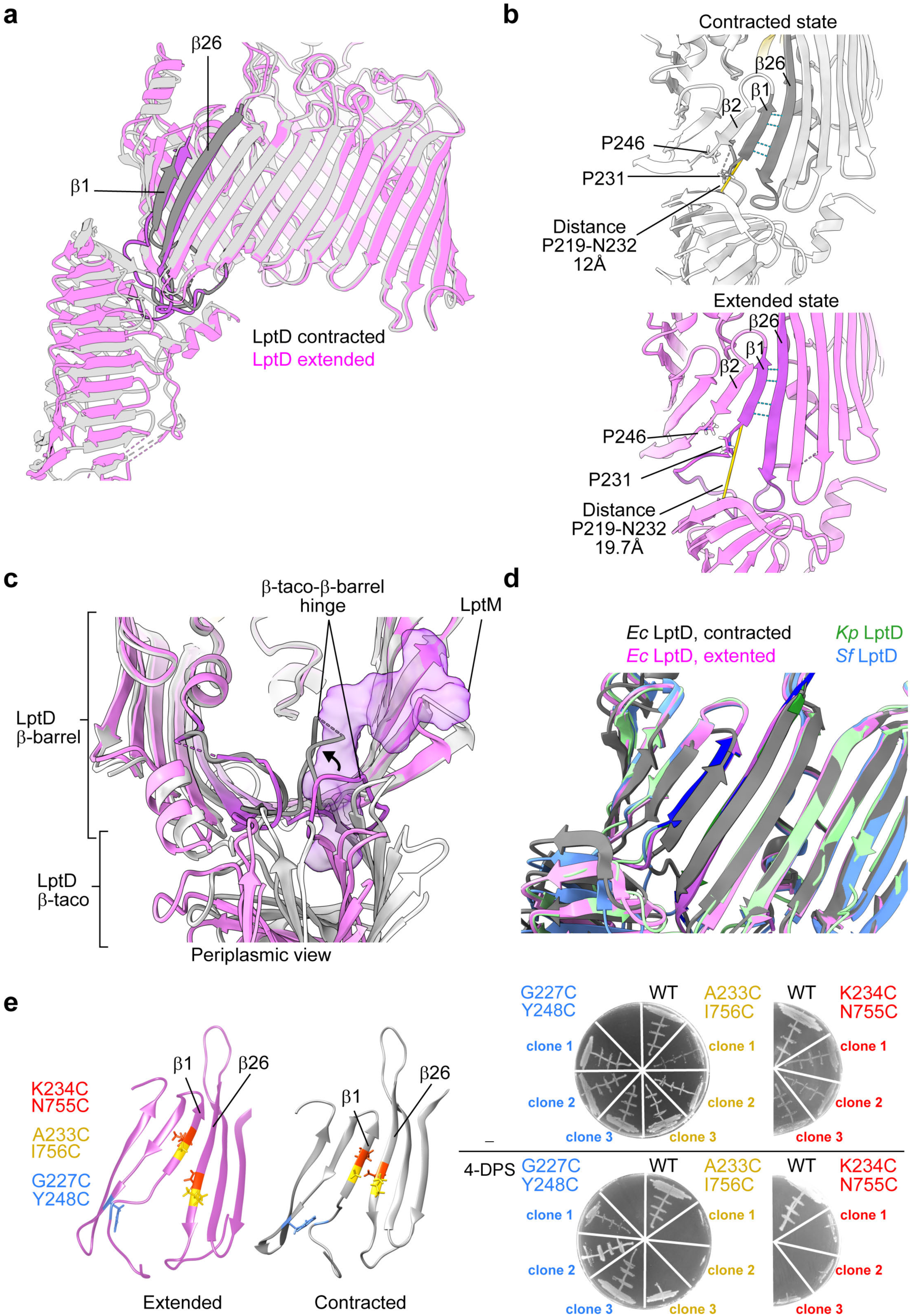
Extended and contracted conformational states of LptD. **a**, Superimposition of LptD conformations observed in the structure of LptDEMY (purple) or LptDEY (grey). The terminal β-barrel strands β1 and β26 are shown in darker colours. Both proteins are shown in ribbon representation. **b**, Zoom on the LptD β-taco/β-barrel interface of LptD in the contracted state (as in the structure of LptDEY) or LptD in the extended state (as in the structure of LptDEMY). The side chains of the pivot residues P246 and P231 are shown. The distance between P219 and N232 is indicated in yellow. **c**, Periplasmic view of the β-taco/β-barrel hinge loop (purple in LptDEMY and grey in the LptDEY) that reorients in the absence of LptM (shown as purple surface). **d**, Superimposition of the structure of LptD in the extended (purple) or contracted (grey) states with the structures of LptD from *Klebsiella pneumoniae* (green) or *Shigella flexneri* (blue). **e**, **Left**, Strands β1-β3 and β24-β26 of the LptD β-barrel domains are shown in ribbon representation, purple in the case of extended LptD (structure of LptDEMY) and grey in the case of contracted LptD (structure of LptDEY). The side chains of the residue pairs replaced with cysteines are shown in different colours. **Right**, Cells deleted of chromosomal *lptD* and harbouring plasmid-borne wild-type or mutant variants of *lptD* with the indicated amino acid substitutions were streaked on LB agar lacking (-) or supplemented with the oxidant 4-DPS. For each *lptD* mutant variant, three clones were selected and tested (see Methods).

### Pivots in the contracted-extended state transition

Whereas high dynamics of the β-taco domains are expected, with different orientations relative to the membrane plane having been previously reported ^24,34,43^, the major structural variations observed in the β-barrel domain of LptD are unprecedented. Two conserved proline residues important for function ^24^, P231 and P246 respectively in LptD β1 and β2, act as pivots for the contracted-to-extended state transitions. P231 serves as a re-orientation point for the luminal loop connecting the β-taco and β1 (Fig. 4b). In the extended state, a portion of this loop prolongs the region of H-bonding between β1 and β2, whereas in the contracted state the loop bends toward the interior of the barrel weakening the β2-β1 pairing (Fig. 4b and c). The other conserved proline, P246, is located where β2 flexes, reducing its tilt angle with the membrane plane at the inner leaflet (Fig. 4b and Extended Data Fig. 11a, Right). This flex point compensates for the increased β-barrel shear number, maintaining the periplasm-oriented hydrophobic belt of the β-barrel within the membrane plane.

Given the identification of these two distinct conformational states of the *E. coli* LPS translocon, we compared our structures to those of LptDE previously determined by X-ray crystallography. We specifically chose LptDE from *K. pneumoniae* (PDB 5IV9) ^24^ and *Shigella flexneri* (PDB 4Q35) ^26^, as these bacteria are closely related to *E. coli*, with a high degree of amino acid sequence identity among their LptD orthologues ^24^. Furthermore, in both cases the structure of full-length LptD was resolved. Our comparison revealed that the pairing pattern of β1-β26 in the extended conformation (LptDEM and LptDEMY) superimposes well with that observed in the LptD structures of *K. pneumoniae* and *S. flexneri* (Fig. 4d and Extended Data Fig. 11b). Furthermore, the recently determined structure of the *E. coli* LPS translocon with bound LptY showed the same β1-β26 pairing pattern observed for *K. pneumoniae* and *S. flexneri* LptD ^41^. Hence the contracted state of LptD identified in our study represents an unprecedented conformation of the LPS translocon.

### Lateral opening of LptD in the contracted state is crucial for function

Our structural analysis uncovered the LptD contracted state in samples lacking LptM. To test whether the LptD contracted state represents a physiological conformation in cells expressing LptM, we employed a cysteine crosslinking approach. A previous study on *Salmonella typhimurium* LptD showed that locking the β1-β26 pairing via inter-strand disulfide bond formation is lethal, supporting a mechanistic model where lateral gate opening is essential for LptD function ^25^. Following this strategy, we tested the effects of forming disulfide bonds between β1 and β26 paired as in the LptD contracted states. As a control experiment, we also tested the effect of forming a disulfide bond between β1 and β2. We generated a diploid *E. coli* strain co-expressing endogenous LptD and a plasmid borne LptD variant with an additional pair of cysteine residues. We then performed deletion of endogenous (wild-type) LptD via P1 phage transduction, and assessed functionality of the mutant LptD variants by streaking the obtained clones on regular media or media containing the oxidant 4,4-dipyridyl disulfide (4-DPS) to facilitates cysteine oxidation (Fig. 4e). The LptD variant harbouring cysteines at position A233C (in β1) and I756C (in β26) or K234C (in β1) and N755C (in β26), which are paired in the contracted state, promoted cell survival only on regular media but not in the presence of the oxidant. In contrast, the LptD variant harbouring cysteines paired in β1 and β2 (G227C and Y248C) promoted cell survival also in the presence of the oxidant (Fig. 4e). This result suggests that, in cells harbouring LptM, the lateral gate β1-β26 paired as in the contracted state can be locked in a closed conformation by an inter-strand disulfide bond, thereby inactivating LptD. We conclude that contracted LptD represents a physiological translocon state and that its lateral opening between β1 and β26 is crucial for function.

### LptM can be docked from the extended state into the contracted conformation

Given the structural differences at the lateral gate between the two LptD states, we questioned whether there was sufficient space to model LptM with LptD in the contracted state. To explore this, we superimposed both conformations and transferred the coordinates of LptM from the extended state to the LptY-bound contracted state of LptDE. This transfer resulted in minimal atomic clashes, which could be further resolved with energy minimization, using the Gromacs MD simulation package ^44^ (see dataset deposited at the Zenodo repository, doi: 10.5281/zenodo.14834225, see also Extended Data Figure 12c, g, i).

### LptM induces gate dynamics, facilitating β1-β26 strand separation

To further assess the impact of LptM and LptY on the core complex, we next performed molecular dynamics simulations of the OM translocon embedded in asymmetric lipid bilayer containing LPS in the external leaflet. The simulations were initiated with LptD in either the extended (LptDEM) or in the contracted (LptDEY) states allowing us to evaluate the contributions of LptM and LptY to translocon lateral gate dynamics (see dataset, doi: 10.5281/zenodo.14834225; Extended Data Figure 12).

We focused our analysis on the pairing of β1-β26 and β2-β1, which vary between the two LptD states. In fact, whereas β1-β26 present alternative pairing patterns, β2-β1 present a number of H-bonds that is higher in the extended conformation as the hinge segment preceding β1 aligns parallel to β2 (Fig. 4a, b and Extended Data Fig 13a). The presence of LptM partially reduced β1-β26 H-bonding, from an average of 3.31 in LptDE and 3.38 in LptDEY to 2.63 in LptDEM and 2.79 in LptDEMY. Interestingly, the addition of KLA built in the β-taco domain of the holo-complex further decreased β1-β26 pairing, averaging only 2.01 H-bonds (Extended Data Fig 13a). Strikingly, when the simulations were initiated with LptD in the contracted state, larger fluctuations in the number of H-bonds were observed not only between β1 and β26 but also between β2 and β1, with both pairs presenting a reduced average of H-bonds in the presence of LptM (Extended Data Fig. 13a). In all cases, β-strand separation was more prominent on their periplasmic side (Extended Data Fig. 13b). Remarkably, full separation of β1-β26 could be observed for a discrete period at least in one of our simulation repeats (Fig. 5a and Supplementary Video). Also in the contracted state, the addition of KLA consolidated the weakening of β-strand pairing (Extended Data Fig. 12a). Taken together, these simulation results indicate that LptM enhances gate dynamics, with this effect being also promoted by the addition of KLA into the β-taco domain. In contrast, LptY had minimal effects on LptD lateral gate dynamics (Extended Data Fig. 13a and b).

**Figure 5.**
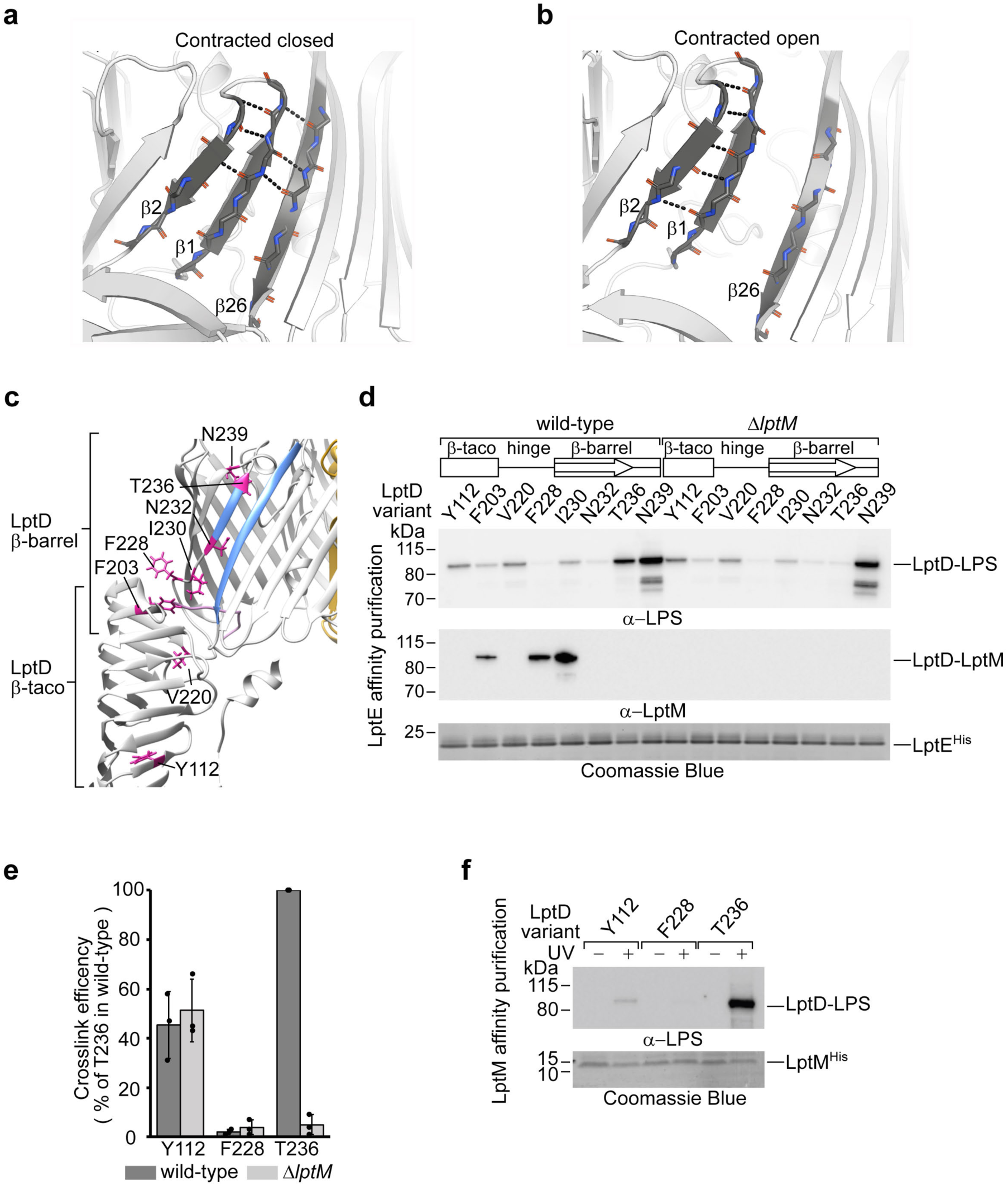
LptM enhances LptD lateral gate dynamics. **a**, Snapshots of the LptD lateral gate conformation in LptDEY containing docked LptM at the start of MD simulations. For clarity, only LptD is shown. β1 and β26 form multiple hydrogen bonds. **b**. Snapshots of the LptD conformation in LptDEY containing docked LptM after 140 ns of MD simulations in one of the three independent runs (Run 1, see Supplementary Video). β1 and β26 are fully separated. In a) and b) β2, β1, and β26 are shown in darker grey. **c**, The structure of LptDEM (extended state) is shown in ribbon representation to highlight the side chains (purple) of LptD amino acids replaced with pBpa for site-directed photocrosslinking. **d**, UV induced photocrosslinking of the indicated strains transformed with pLptDE^His^ harbouring amber mutations at the indicated amino acid positions of LptD. Upon in vivo photocrosslinking, the envelope fractions were solubilized with DDM and subjected to nickel-affinity purification of LptE^His^. **e**, Quantitation of ECL signals of LptD-LPS crosslinking adducts for the indicated LptD positions in cells expressing or lacking LptM, as shown in d). Data are represented as means, ± s.e.m. (*n* = 3). Values are normalized to the crosslink intensity determined for LptD containing pBpa at position T236 expressed in wild-type cells. **f**, UV induced photocrosslinking of the indicated strains transformed with pLptDEM^His^ harbouring amber mutations at the indicated positions in LptD. Solubilized membranes were prepared as in d), and subjected to nickel-affinity purification of LptM^His^.

### LptM operates at the translocon during LPS gating

Given its positioning at the β-taco/β-barrel interface and its effect on lateral gate dynamics, we investigated whether LptM influences the interaction between distinct domains of LptD and LPS. LptM is required for the proper oxidative maturation of LptD. Nevertheless, upon overproduction of LptDE in Δ*lptM* cells, we showed that the majority of LptD (approximately 70%), is correctly oxidized. This fraction increases to 95% when LptM is overproduced together with LptD and LptE ^36^. Despite this marginal difference in LptD oxidation, we probed distinct LptD domains to monitor their interactions with LPS. We introduced the photoactivable amino acid analogue para-benzoyl phenylalanine (pBpa) at specific positions within the β-taco domain, the hinge region and β1 (Fig. 5c). We aimed to maximize the probability of detecting interactions between LPS and translocon intermediates by replacing with pBpa the amino acids whose side chains project into the hydrophobic cavity of the β-taco or toward the internal lumen of the β-barrel. Upon UV irradiation and translocon purification, we detected crosslinks between LptD and LPS along the β-taco domain and β1 (Fig. 5d). Notably, positions T236 at the end of β1 and N239 in the β1-β2 loop showed the highest crosslinking efficiency. The side chains of T236 and N239 remain oriented toward the interior of the barrel during our MD simulations, although that of T236 showed a higher degree of dihedral angle variation (Extended Data Fig. 14a and b). The efficient interaction of these amino acid positions with LPS indicates that the apical region of β1, which repositions during the contracted-extended conformational switch observed in our structural analyses, represents a major LPS binding site. Positions near the β-taco/β-barrel interface interact with LPS with lower efficiency, and these were also found to bind LptM, consistent with our structural analysis. Crosslinking to LPS within the β-taco domain occurred with a similar efficiency regardless of whether LptM was produced. Remarkably, at position T236 in the C-terminal end of β1, the efficiency of LPS-crosslinking was drastically reduced by >90% in the absence of LptM (Fig. 5d and e). Taken together, these results suggest that LptM influences the mature, active translocon, facilitating LPS access to a position within the β-barrel lateral gate.

Finally, we investigated whether LptM plays a direct role at the active translocon during LPS secretion. Using a similar site-directed crosslinking approach, we selected three positions within LptD to monitor interactions with LPS by LptM^His^ pull-down, thereby assessing for the simultaneous binding of LPS and LptM to the translocon. Our analysis clearly shows that LPS binds to position 236 of LptD without requiring the release of LptM from the translocon (Fig. 5f). Notably, in both the contracted and the extended states of LptD, position 236 is closer to the external leaflet compared to the N-terminus of LptM near the lateral gate, indicating that LPS bypasses LptM during translocation.

## DISCUSSION

It has been unknown how the LptD β-barrel can open laterally to release LPS into the external leaflet of the OM. High resolution structures of LptD have shown extensive pairing between LptD β1 and β26, implying a high energetic barrier to lateral gate opening^24–26,34^. In our study, we demonstrate an unprecedented conformation of the LptD β-barrel that can open via a mechanism facilitated by LptM. We uncover a conformational switch of the LptD lateral gate explaining how LPS is directed to the membrane external leaflet.

We have previously identified LptM as a translocon component that is assembled with an LptD folding intermediate at the BAM complex. Predicted to bind near the β-taco/β-barrel interface, we showed that LptM promotes the subsequent step of LptD oxidative maturation characterized by the formation of inter-domain disulfide bonds. Given that LptM occupies a portion of the translocon near the predicted path of LPS transport, with this study we aimed at addressing how LptM influences translocon activity. Our biochemical and structural analyses identified a further translocon component, LptY, which binds the LptD β-taco domain externally to its LPS binding groove, as shown also by a recent study ^41^. Differently from LptM, LptY does not appear to influence LptD oxidative maturation. We showed that together LptM and LptY stabilize the β-taco domain thereby facilitating the connection of the OM translocon to the IM components of the Lpt machinery (Fig. 6a). It remains to be determined whether the binding of LptY to LptD is regulated *in vivo*. Given that LptY presents a lipocalin-like fold and possesses two Cys residues predicted to form a disulfide bond, it is possible that a ligand or the periplasm oxidative state can influence the binding of LptY to LptD.

**Figure 6.**
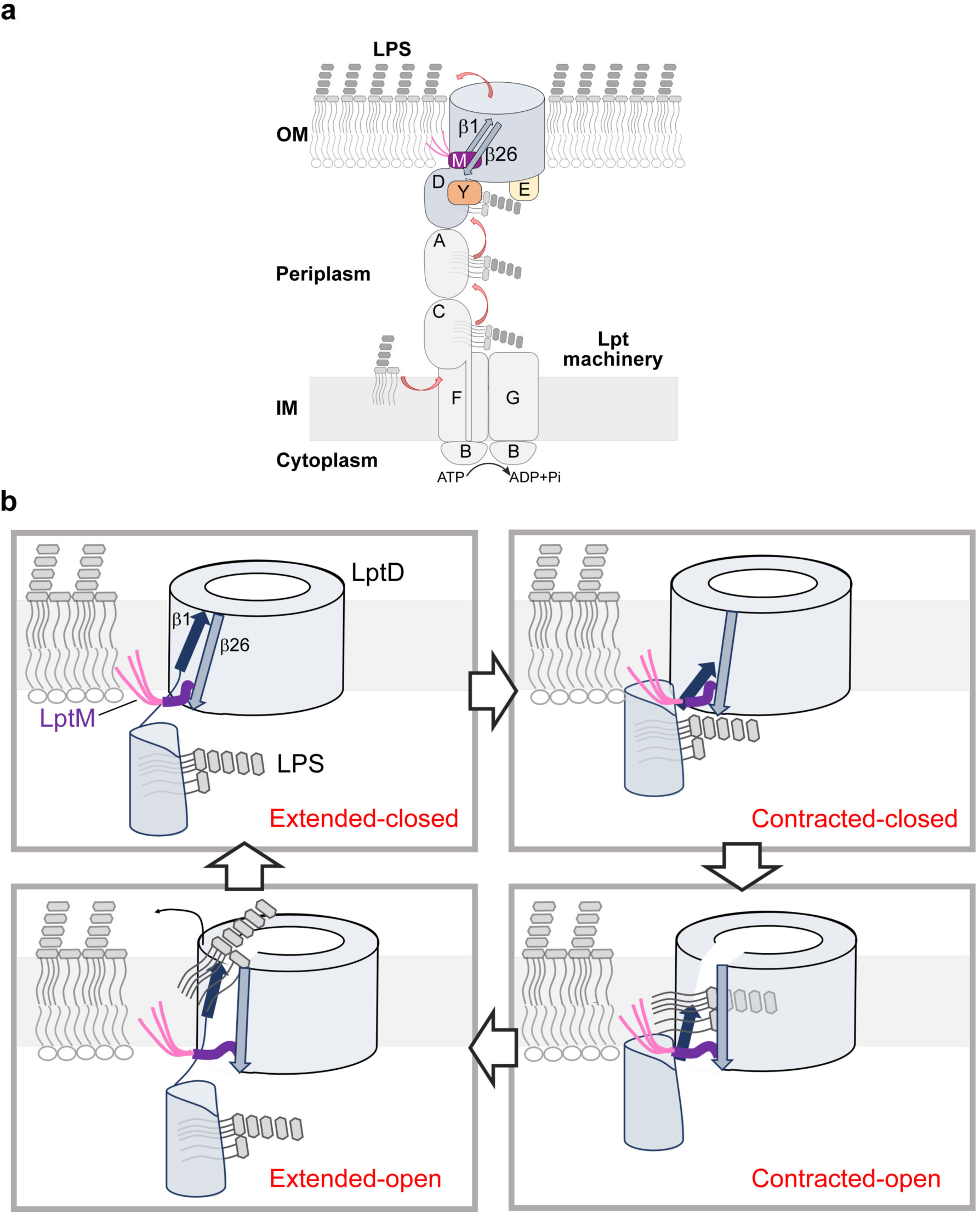
LptD operates by switching between contracted and extended conformational states. **a**, Diagram of the Lpt pathway including the subunits LptM and LptY. **b**, Mechanism of LPS insertion proposed on the basis of LptD conformational dynamics and functional assays described in this study. **Top left**, LPS binds the β-taco domain of LptD (extended-closed state). **Top right**, LptD contracts, bringing LPS close to β1 of the LptD β-barrel domain (contracted-closed state). **Bottom right**, LptM facilitates lateral opening of LptD in the contracted state (contracted-open state); the LPS sugars enter the lumen of LptD β-barrel, whereas its lipid tails move from the taco into the OM bilayer, bypassing LptM. **Bottom left**, LPS moves together with β1 towards the OM external leaflet, which restores the LptD extended state with an open lateral gate (extended-open state). LPS is then ejected from the LptD β-barrel domain, resetting LptD in the extended-closed state. For clarity, LptE and LptY are not shown.

Remarkably, by stabilizing the LptD β-taco domain, LptY binding in the absence of LptM enabled the identification of a contracted LptD state, in which the β-taco domain and β1 of the β-barrel domain have converged, thereby contracting their interface. Indeed, in the contracted state β1 has slid along β26 of two amino acid positions toward the periplasm, revealing an unprecedented rearrangement of the lateral gate β-strands that provides direct evidence for their unzipping. Although we observed the contracted state in the absence of LptM, our functional assays with a LptD variant harbouring crosslinkable Cys residues in β1 and β26 demonstrates the relevance of the contracted state for LptD function also in the presence of LptM. MD simulations of contracted LptD with modelled LptM showed enhanced lateral gate dynamics, which might explain why our cryo-EM structural analysis revealed the contracted LptD state only in the absence of LptM.

The LptM-induced lateral gate dynamics are likely to influence LPS transport. Our crosslinking results show that LptM remains associated with the translocon when LPS crosses the LptD lateral gate. The first 10 N-terminal residues of LptM interact with the internal wall of LptD β1-β4, whereas the C-terminal moiety of LptM was not structurally resolved, suggesting that it is highly dynamic. We speculate that the C-terminal moiety of LptM can occupy the lumen of the barrel, as previously predicted by AlphaFold 2 ^36^ but also move out of the translocon, *e.g.* by retrograding toward the periplasm. At the active translocon, displacement of LptM C-terminal moiety would free the β-barrel lumen for access by LPS. Indeed, we identified luminal oriented residues of LptD β1 and of the β1-β2 loop that can be crosslinked to LPS with strong efficiency. An interaction at the apical region of β1 at the lateral gate slit is strongly affected by the lack of LptM, corroborating the role of this translocon component in functionalizing the LptD lateral gate.

Based on our structural and functional characterization of contracted LptD, we propose that the translocon operates by alternating between contracted and extended states. Contraction of LptD brings β1 of the β-barrel closer to the β-taco domain, which can facilitate LPS transfer to the LptD lateral gate (Fig. 6b, Contracted-closed). In the contracted state, LptM enhances LptD lateral gate opening, thereby facilitating the insertion of the LPS acyl tails into the membrane and entry of its saccharides into the β-barrel lumen (Contracted-open). This step also requires that LPS crosses the luminal gate made of the β-taco/β-barrel hinge loop and a C-terminal segment of LptD ^32^. We hypothesize that, at this stage, the acyl tails of LptM might help prevent the phospholipids of the OM inner leaflet from accessing the open lateral gate. By interacting with LPS and by repositioning during the contracted-to-extended state transition, β1 performs a stroke-like movement that can push LPS toward the external leaflet of the OM (Extended-open). Intriguingly, the connection of LptD to LptB_2_FG via stacked β-taco domains suggests that ATP hydrolysis and LPS extraction at the IM could contribute to power the β-taco/β-barrel contraction/extension mechanism resulting in the stroke movement of β1 at the OM. With LptD in the extended-open state, repulsion of LPS phosphate groups by negatively charged residues in the translocon lumen ^24,34^ aids in the ejection of LPS through the lateral gate. The LPS release step can be further facilitated by LptE ^45^ and by interactions of LPS with divalent cations in the external leaflet. The translocon would then be ready to reset for the transport of a new molecule of LPS (Extended-closed).

The essential role of the Lpt machinery and its partial exposure to the cell surface make it an ideal target for the design of new antibiotics ^46–48^. A detailed understanding of its components and their architecture to form a functional Lpt transenvelope machinery is of fundamental importance for drug-targeting. The new Lpt components characterized in this study are both highly conserved in pathogenic species of *Enterobacteriaceae* and *Erwiniaceae*. In particular, LptM is also present in the threatening pathogens of the *Yersinia* and *Serratia* genera. Most importantly, the demonstration that LptD operates by alternating between contracted and extended states establishes a detailed framework illustrating how LPS is assembled at the surface of the OM. The improved understanding of the LPS translocon holo-complex has major implications for antibiotic development.

## METHODS

See Supplementary Information

## ACKNOWLEDGMENTS

Research in R.I.’s laboratory was funded by the Agence Nationale de la Recherche (ANR-23-CE11-0025-01), the China Scholarship Council fellowships to H.C., the FRM PhD fellowships to Y.A.A. and the FRM Fourth-Year PhD Fellowship to H.C. Cryo-EM imaging of samples was sponsored by the iNEXT Discovery program (PID: 22756) to R.I. Research in R.F.’s laboratory was funded by the Agence Nationale de la Recherche (ANR-21-CE44-0002) and the CNRS. We acknowledge the METi imaging facility, member of the national infrastructure France-BioImaging supported by the Agence Nationale de la Recherche (ANR-10-INBS-04). Research in J.M. laboratory was supported by the Agence Nationale de la Recherche: ProFI projects (ANR-10-INBS-08 & ANR-24-INBS-0015). P.J.S. acknowledges the NIH (R01AI174416 (PI: M. Stephen Trent)), Wellcome (208361/Z/17/Z), MRC, BBSRC, EPSRC and the Howard Dalton Centre for funding. P.J.S. and D.G. acknowledge Sulis at HPC Midlands+, which was funded by the EPSRC on grant EP/T022108/1, and the University of Warwick Scientific Computing Research Technology Platform for computational access. This project made use of time on ARCHER2 granted via the UK High-End Computing Consortium for Biomolecular Simulation, HECBioSim (http://www.hecbiosim.ac.uk), supported by EPSRC (grant no. EP/R029407/1).

## DATA AVAILABILITY

LptDE model information is available under accession PDB 9I9Z and the density map under accession EMDB 52773. LptDEY map is available under EMDB 52777 and the pertaining LptDE model under accession PDB 9IA0. LptDEM model information is available under accession PDB 9IA2 and the density map under accession EMDB 52778. LptDEMY map is available under EMDB 52779 and the pertaining LptDEM model under accession PDB 9IA5. MD simulations data are available at the Zenodo repository, doi: 10.5281/zenodo.14834225.

## AUTHOR CONTRIBUTIONS

H.C., V.M. performed genetic analyses, protein purifications and functional assays, with contributions from Y.A.A. and A.C.-S.; S.B. prepared cryo-EM grids; R.F and A.S. determined the cryo-EM structures; P.J.S. performed structural modeling and MD simulations, and analyzed the data together with D.G.; Y.Q. performed phylogenetic analysis; J.M. performed native-MS; J.M., M.M. and C.F. performed LC-MS/MS analysis; H.C., A.S., V.M., Y.Q., J.M., D.G. prepared the figures; R.I., R.F. and P.J.S. wrote the manuscript with contributions from J.M., Y.Q. and V.M. and A.C.-S.; R.I. conceived the research. R.I., R.F. and P.J.S. supervised the work and acquired funding. All authors discussed the results and commented on the manuscript.

## EXTENDED DATA FIGURES AND LEGENDS

**Extended Data Fig. 1.**
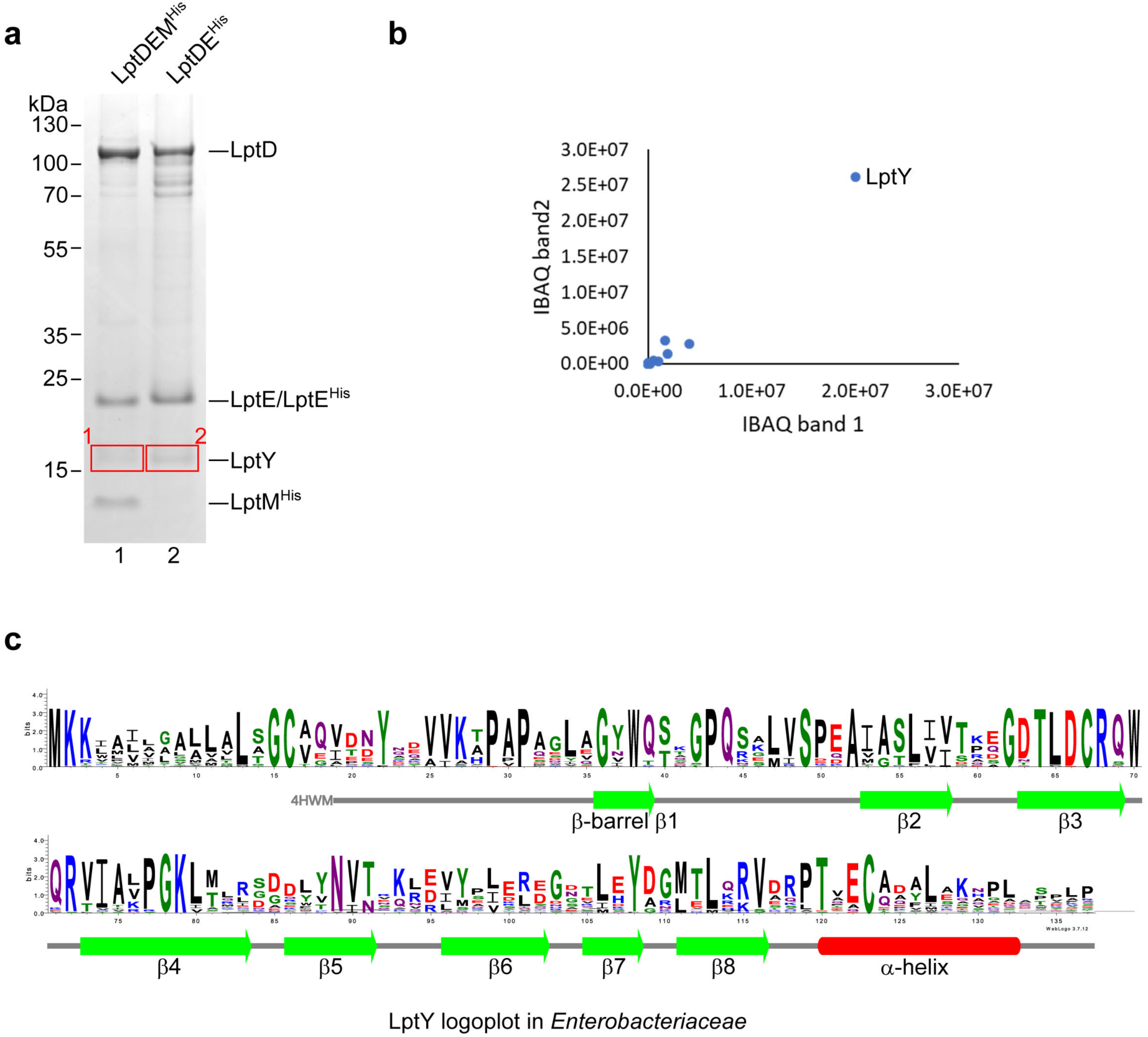
SDS-PAGE based, label-free bottom-up proteomic analysis of LPS translocon complexes. **a**, LptDEM^His^ and LptDE^His^ samples, prepared as in Fig. 1b, were separated on SDS PAGE. Gel bands running with an apparent molecular weight of approximately 15 kDa (red boxes) were excised and their protein contents were analysed by bottom-up mass-spectrometry. **b**, The plot represents iBAQs (intensity-based absolute quantification) of the excised band 1 (iBAQ1, LptDEM^His^) vs. excised band 2 (iBAQ2, LptDE^His^). The iBAQ value is obtained by dividing protein intensities by the number of theoretically observable tryptic peptides (see Methods). This analysis clearly identifies LptY as the main protein present in the excised band, in both LptDEM^His^ and LptDE^His^ purified complexes. **c**, Logoplot of LptY amino acid sequence. The logoplot is based on the alignment of the LptY sequences of the genus reference genomes of *Enterobacteriaceae*.

**Extended Data Fig. 2.**
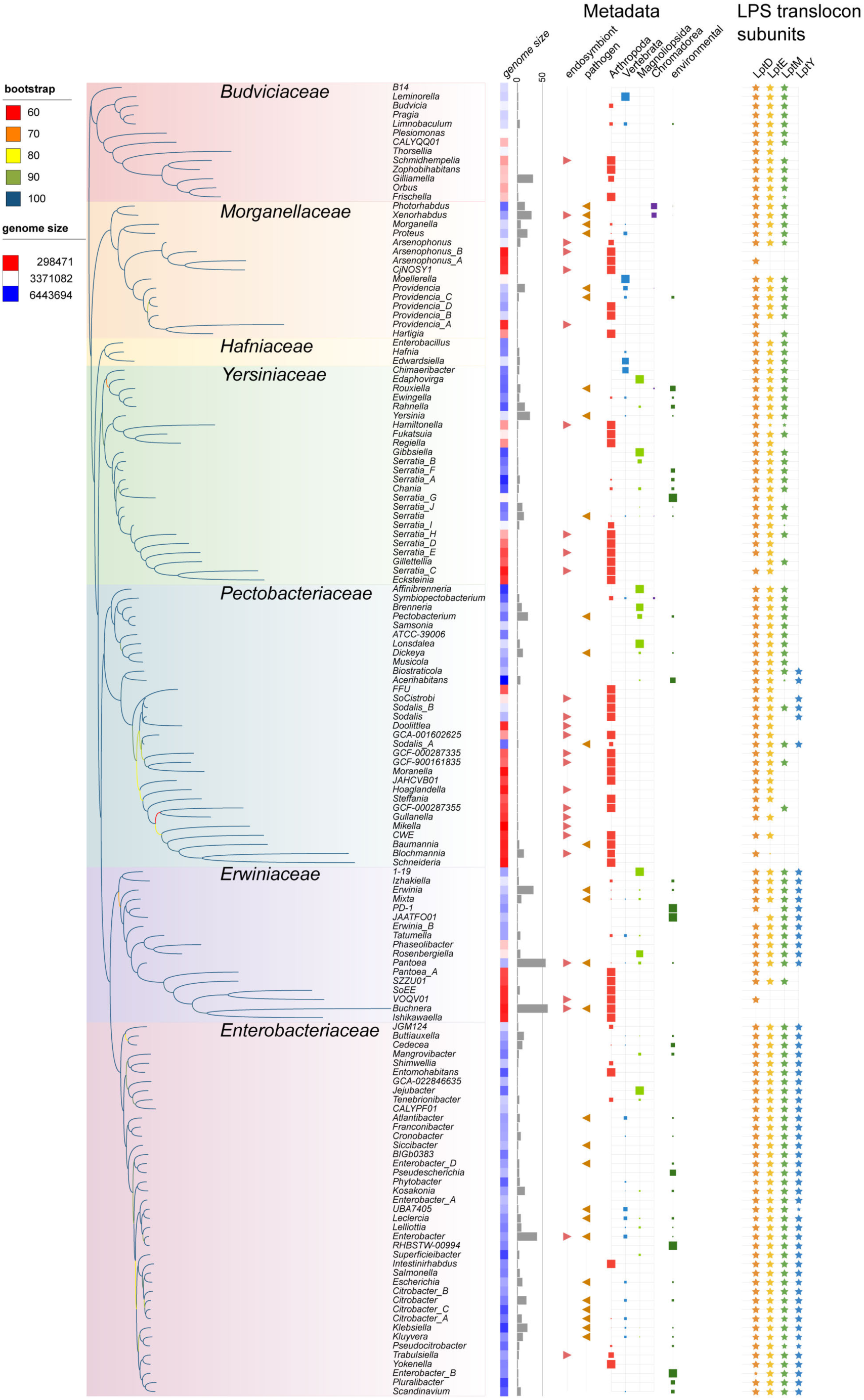
Distribution of LptY proteins in *Enterobacterales* in relation to genome features and the LPS transporter. We used a single representative genome per genus of *Enterobacterales*. The phylogenetic tree inferred with *IQ-TREE2* (Minh et al. 2020) on an alignment of 145 concatenated sequences and 80417 sites. The lengths of the branches are proportional to the distances inferred by *IQ-TREE2*. The branch supports, reported as branch color gradient, were assessed with the SH-like approximate likelihood ratio test (-alrt 1000). The tree was rooted with *Budviciaceae* family. *Enterobacterales* are divided in seven families. Columns annotations, genome size: colour gradient of the genome sizes with small genomes in red and large genomes in blue, grey histogram: number of genomes in each genus, Metadata: metadata extracted from NCBI’s BioSample database, LPS translocon holo-complex: LptDEMY proteins. Intra-genus variability is illustrated by the size of the symbol, which is proportional to the frequency with which the feature is observed in the genomes of that genus. The figure has been generated using iTOL (see Methods).

**Extended Data Fig. 3.**
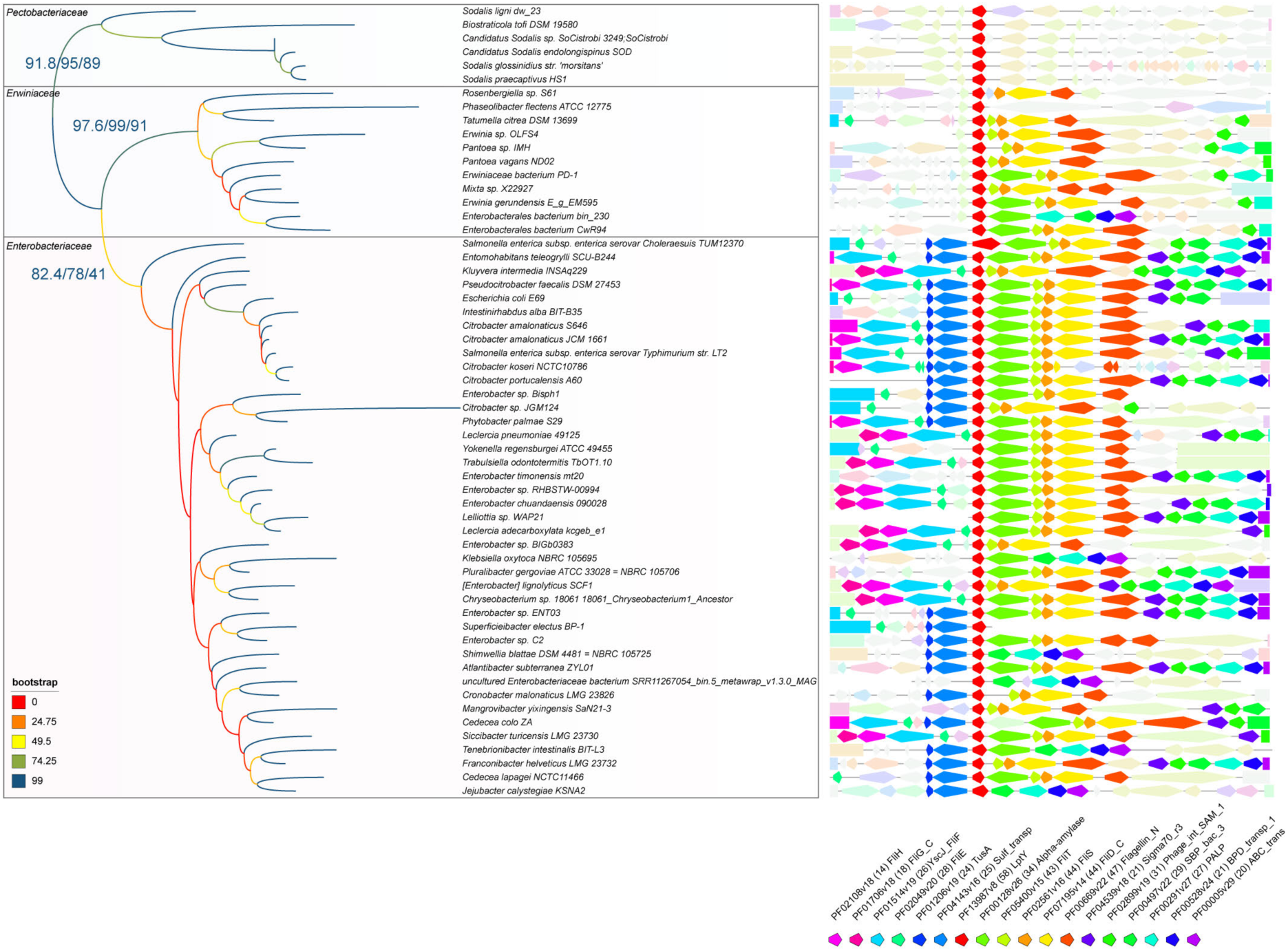
LptY protein tree in the *Enterobacterales* genera. The tree was inferred with *IQ-TREE2* (see Methods). Branch supports were estimated using the ultra-fast bootstrap approximation, the SH approximate likelihood ratio test and the non-parametric bootstrap. The values obtained were displayed on the three deepest branches of the tree and the non-parametric bootstrap were plotted on the tree as a gradient of branch color. The division of *Enterobacterales* into three families was reported. The genomic contexts of the *lptY* genes extracted 5000 nucleotides upstream and 10000 nucleotides downstream of the *lptY* genes were displayed. A colour code was used to highlight genes sharing the same Pfam domain annotation. Domain accession, name and frequency are indicated. Very light color were used for genes present in fewer than 14 neighbourhoods. The figure has been generated using iTOL (see Methods).

**Extended Data Fig. 4.**
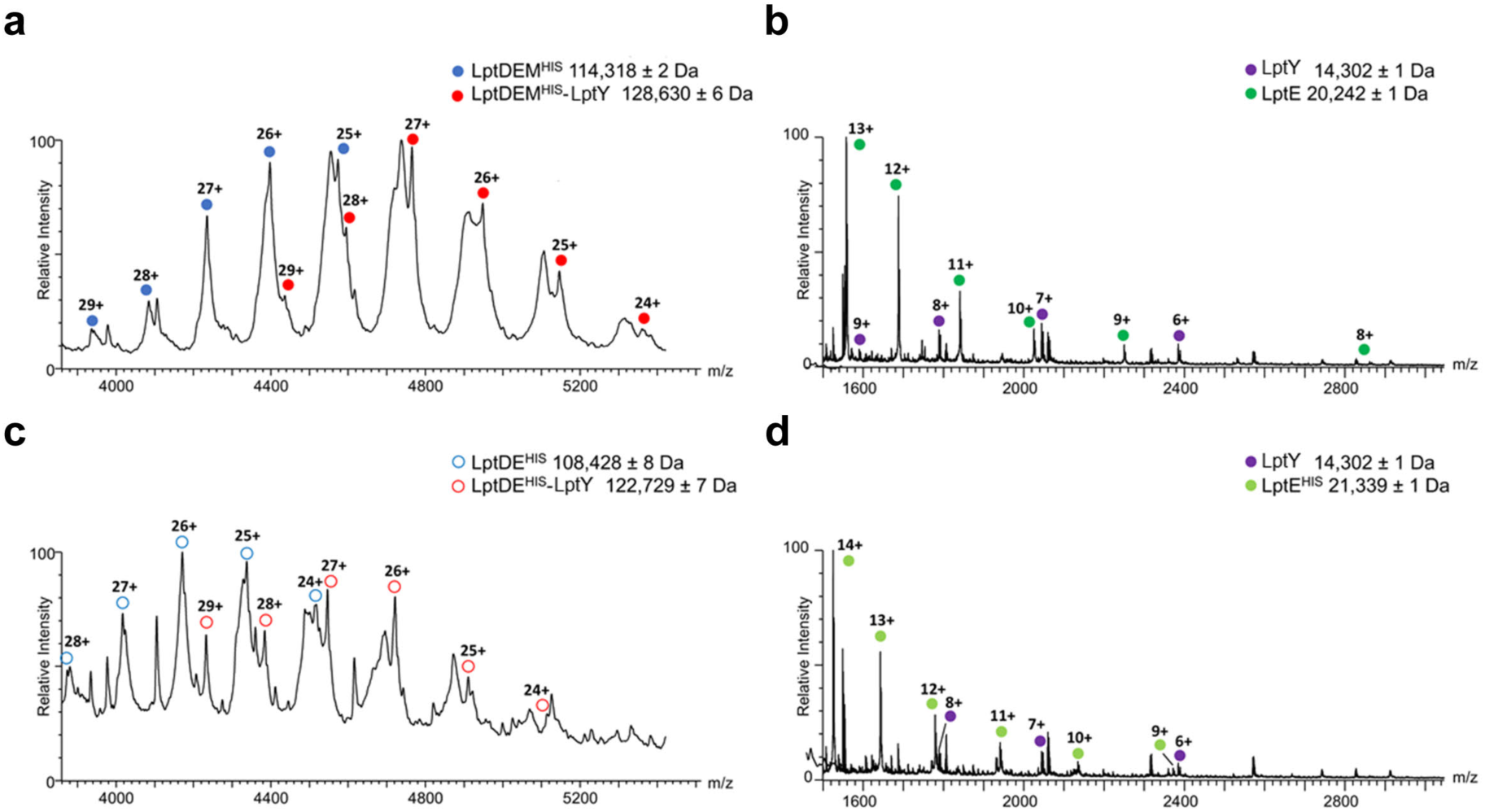
Native MS of the LPS translocons. Mass spectra of (**a**) LptDEM^His^ and (**b**) LptDE^His^ aquired under conditions to maintain non-covalent interactions shows the presence of the corresponding translocons alone and with a mass shift of of 14,3 kDa. A gas-phase dissociated monomer with MW=14,302 Da, corresponding to LptY can be observed in the lower m/z region for both the (**c**) LptDEM^His^ and (**d**) LptDE^His^ complexes.

**Extended Data Fig. 5.**
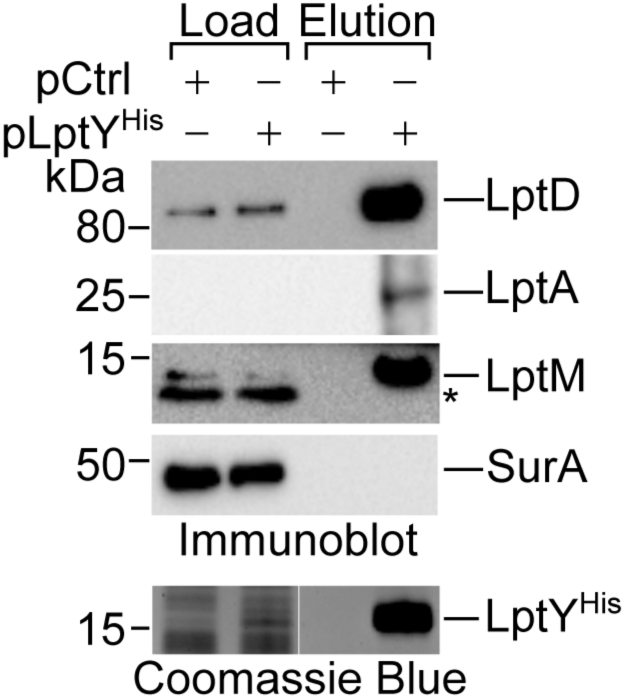
Pull down of LptY co-isolates components of the LPS OM translocon and LptA. Δ*lptY* transformed with the empty vector pCtrl or with pLptY^His^ were subjected to envelope fractionation, solubilization with DDM and nickel-affinity purification of LptY^His^. Imidazole eluted proteins were subjected to SDS-PAGE and revealed by immunoblotting using the indicated antisera or Coomassie Blue staining to visualize the bait protein. Load: 0.5%; Elution 100%; * indicates a non-specific reaction.

**Extended Data Fig. 6.**
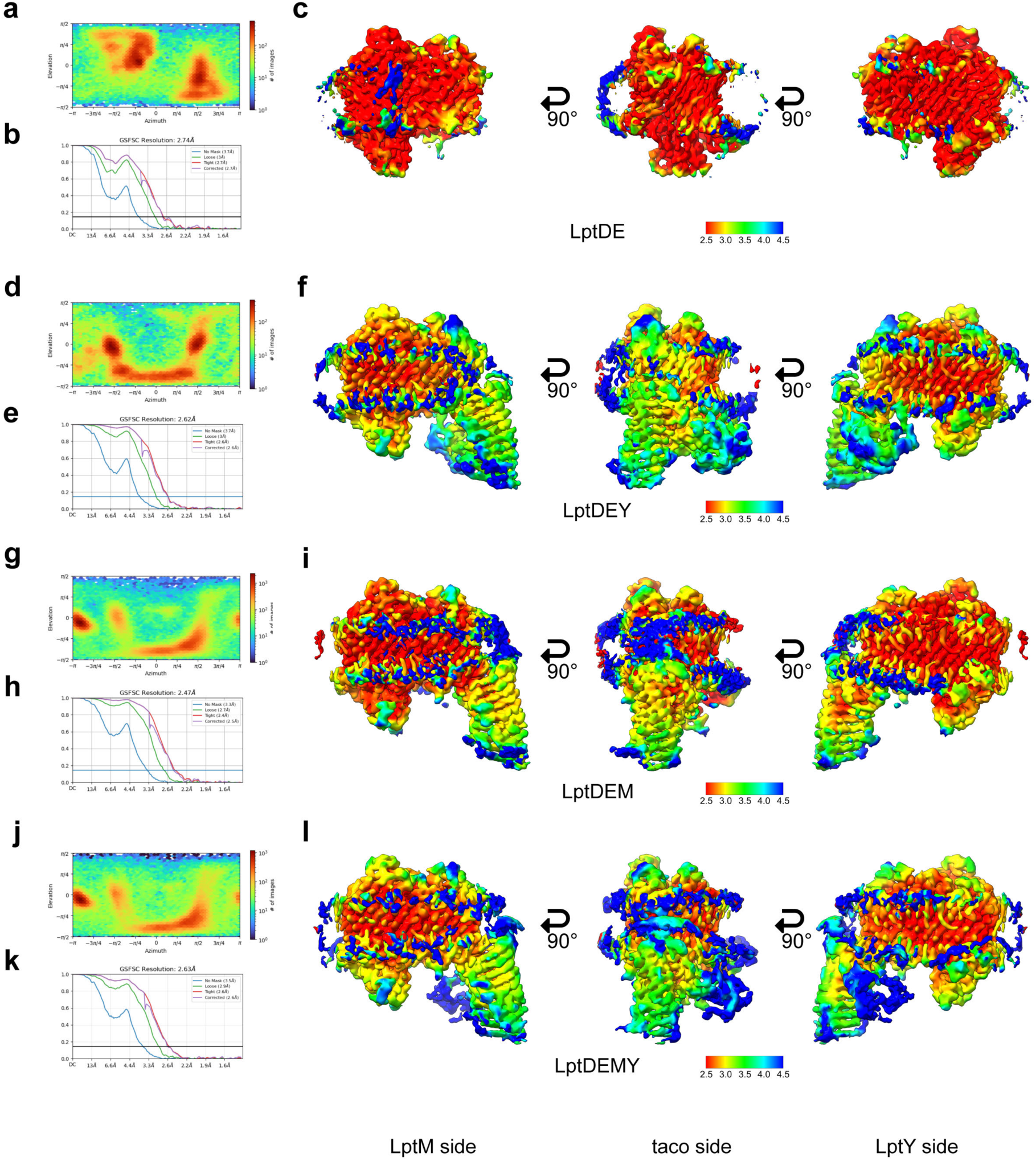
Local refinement of the LptDE heterocomplexes. **a**, Particle orientation distribution of the LptDE heterodimer, obtained in Cryosparc. **b**, Fourrier Shell Correlation (FSC) map obtained from half-maps in Cryosparc. **c**, Local-filtered map of LptDE, at level 0.219, coloured by local resolution. **d**, Particle orientation distribution of the LptDE-LptY oligomer, obtained in Cryosparc. **e**, Fourrier Shell Correlation (FSC) map obtained from half-maps in Cryosparc. **f**, Local-filtered map of LptDEY, at level 0.168, coloured by local resolution. **g**, Particle orientation distribution of the LptDEM complex, obtained in Cryosparc. **h**, Fourrier Shell Correlation (FSC) map obtained from half-maps in Cryosparc. **i**, Local-filtered map of LptDEM, at level 0.161, coloured by local resolution. **j**, Particle orientation distribution of the LptDEMY complex, obtained in Cryosparc. **k**, Fourrier Shell Correlation (FSC) map obtained from half-maps in Cryosparc. **l**, Local-filtered map of LptDEMY, at level 0.171, coloured by local resolution.

**Extended Data Fig. 7.**
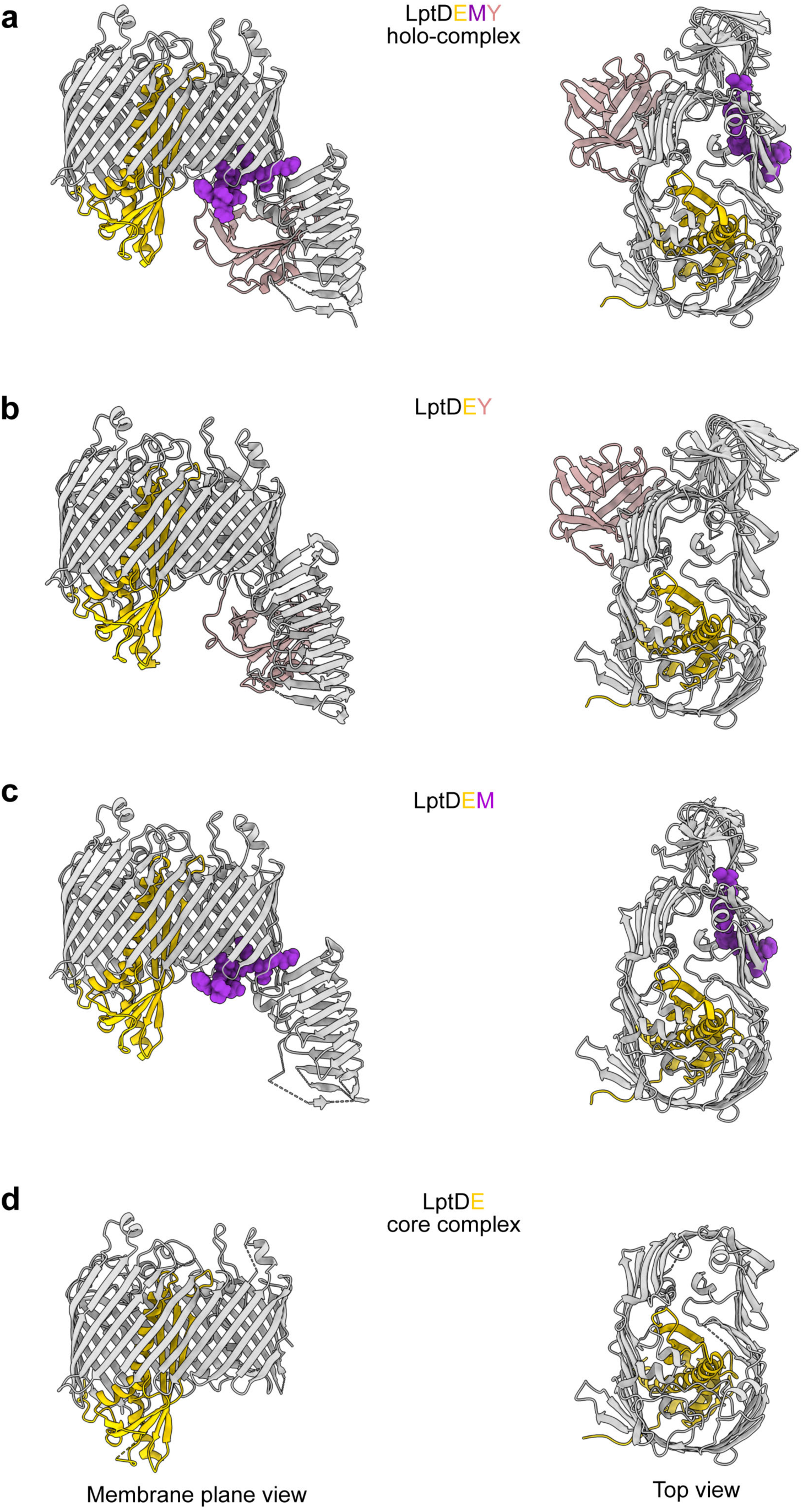
Views of the resolved structure. Ribbon representations of the resolved structures seen from the membrane plane (**letf**) and from the top (**right**) with LptD in grey, LptE in yellow, LptM in purple and LptY in pink. **a**, LptDEMY holo-complex **b,** LptDEY complex. **c**, LptDEM complex. **d**, LptDE core complex.

**Extended Data Fig. 8.**
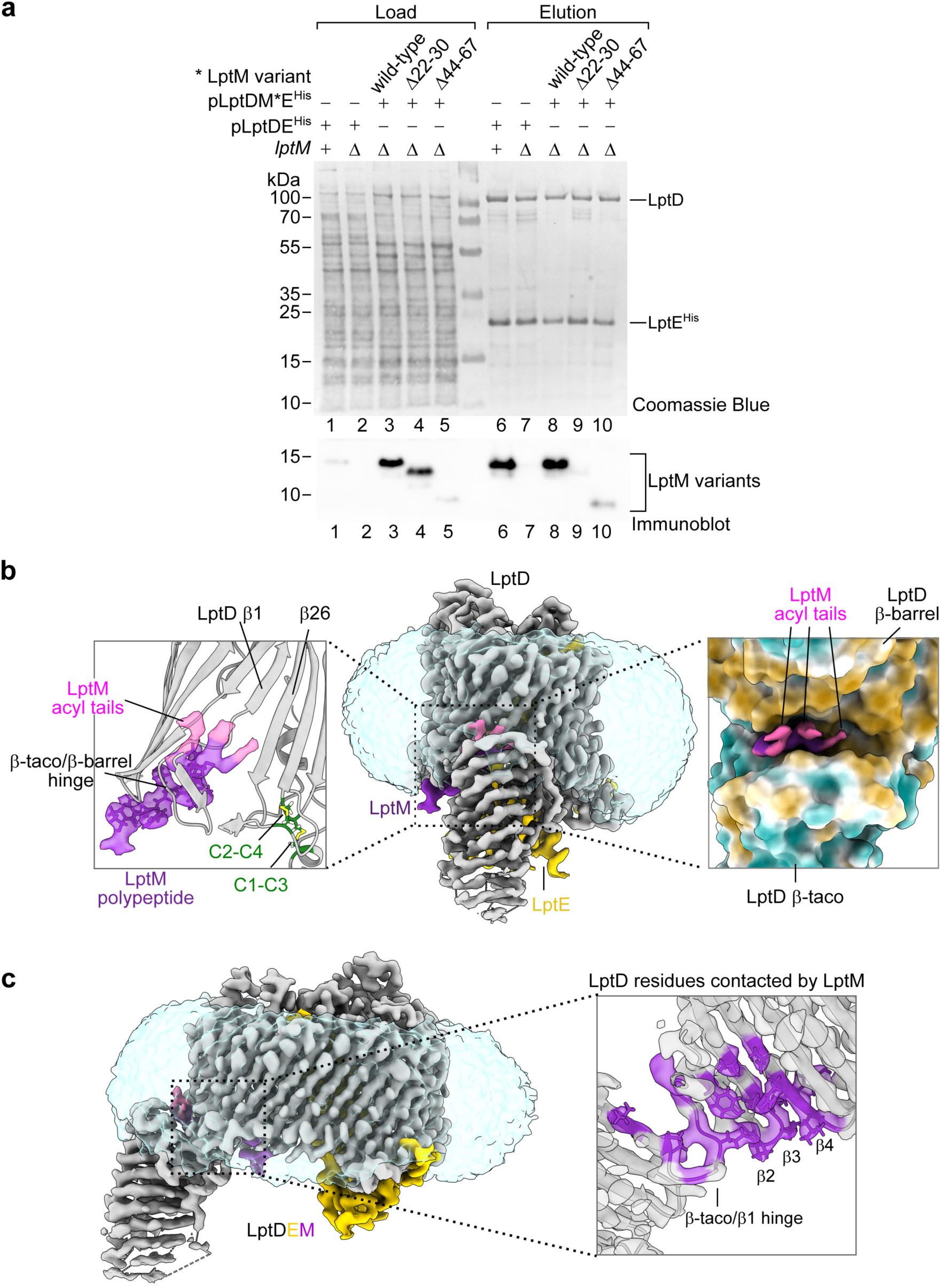
The N-terminal portion of LptM interacts with the translocon. **a**, Wild-type or Δ*lptM* cells transformed with the indicated plasmids were subjected to envelope fractionation, solubilization with a mild detergent and nickel-affinity purification of LptE^His^. Only the LptM variants containing the protein N-terminal portion (wild-type LptM and LptM^1144–67^) were enriched in the elution fractions. Load: 1%; Elution 100%. **b, Left**, Zoom on LptM in the structure of LptDEM complex, highlighting the electron density corresponding to the acyl tails of LptM (pink) and the electron density corresponding to LptM N-terminal moiety (purple) situated near the lateral gate of LptD (β1-β26) and between the β-taco/ β-barrel hinge and the LptD inter-domain disulfide bonds. LptD Cys31 (C1), Cys173 (C2), Cys724 (C3) and Cys725 (C4) are shown in green with disulfide bonds in yellow. **Central**, cryo-EM map of the LptDEM complex coloured according to the protein chains (LptD in grey, LptE in yellow, LptM in purple). **Right**, zoom on the lipid tails of LptM and the surrounding surface of LptD coloured based on hydrophobicity of the amino acid side chains (brown, hydrophobic; cyan, hydrophilic). **c, Left**, cryo-EM map of the LptDEM complex coloured according to the protein chain (LptD in grey, LptE in yellow, LptM in purple). **Right**, internal section of the LptD β-taco/β-barrel hinge region. The LptD residues in contact with LptM are shown in purple.

**Extended Data Fig. 9.**
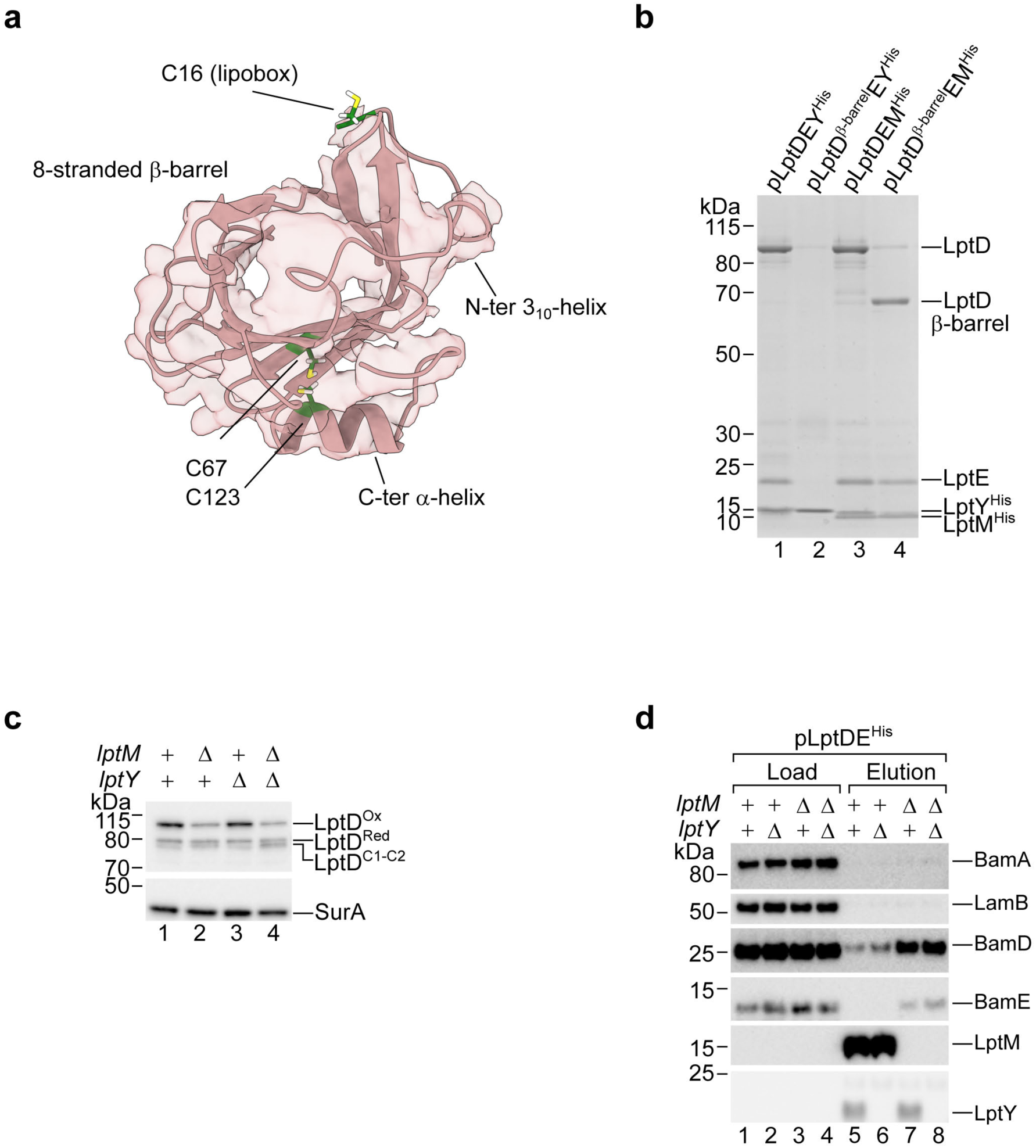
Characterization of LptY interaction with LptD. **a,** The AlphaFold 2-predicted model of LptY (shown in pink ribbon representation) is docked into its corresponding cryo-EM density map shown as a transparent surface. The main structural features of LptY are indicated. The lipobox cysteine (C16) that anchors LptY to the membrane, and the conserved cysteines (C67 and C123) that form an intramolecular disulfide bridge are highlighted as green sticks. **b**, Wild-type cells transformed with the indicated plasmids were subjected to envelope fractionation, solubilization with DDM followed by nickel-affinity chromatography. Proteins in the elution fractions were separated by SDS-PAGE and stained with Coomassie Blue. **c**, Total cell lysates of the indicated strains were subjected to non-reducing SDS-PAGE and immunoblotting using the indicated antisera. **d**, The indicated strains transformed with pLptDE^His^ were subjected to envelope fractionation, solubilization with DDM followed by nickel-affinity purification of LptE^His^. Load and elution fractions were subjected to SDS-PAGE and immunoblotting using the indicated antisera. Load: 0.1%; Elution 100%

**Extended Data Fig. 10.**
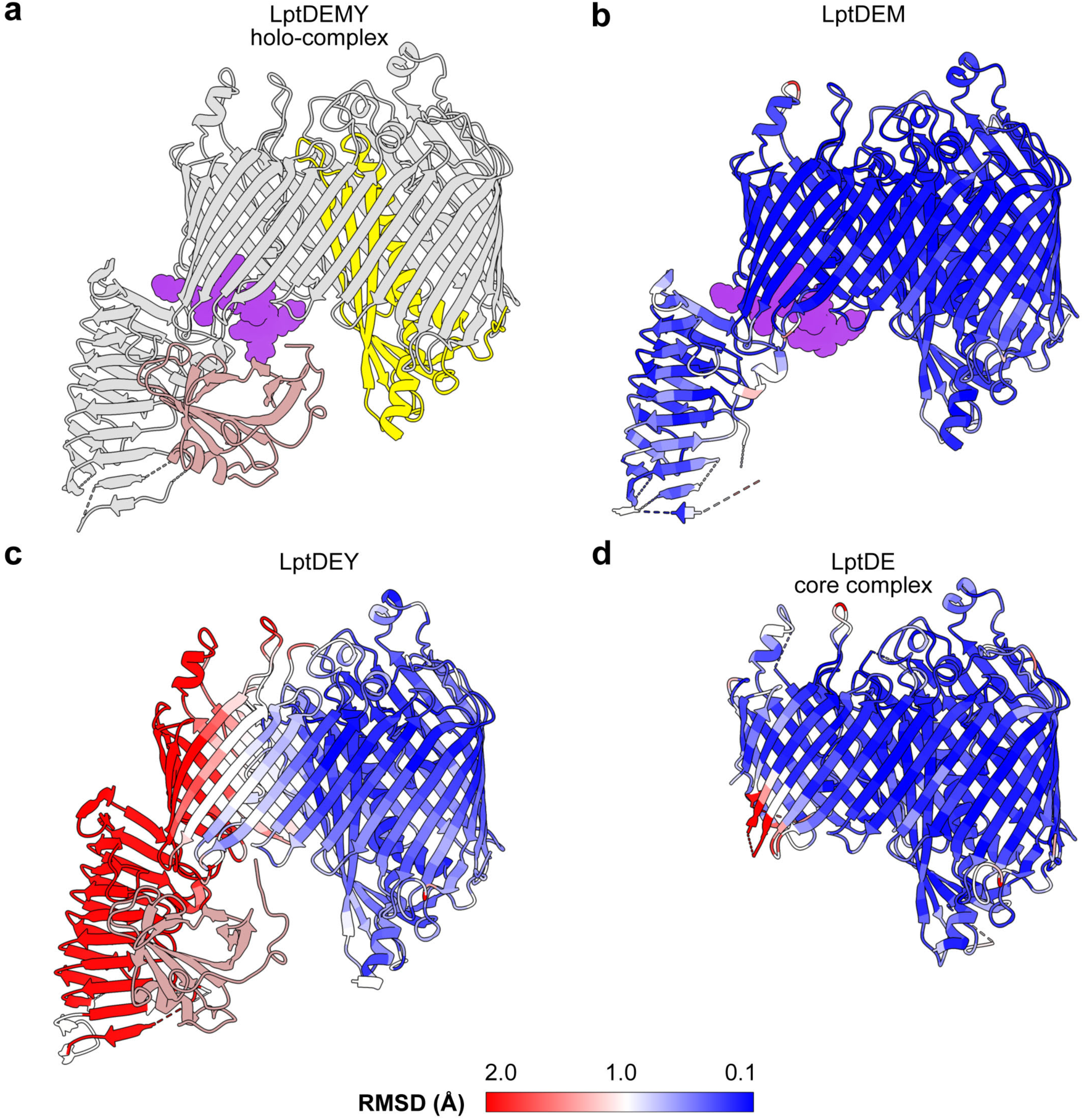
Structural comparison of translocon complexes. Root mean square deviation (RMSD) of the superimposed structures are shown for LptD and LptE: **a**, LptDEMY holo-complex is used as a common reference to calculate the RMSD values. **b**, LptDEM versus LptDEMY. **c**, LptDEY versus LptDEMY. **d**, LptDE core complex versus LptDEMY. b-d, RMSD data are shown on a blue to red scale and coloured onto a protein ribbon representation.

**Extended Data Fig. 11.**
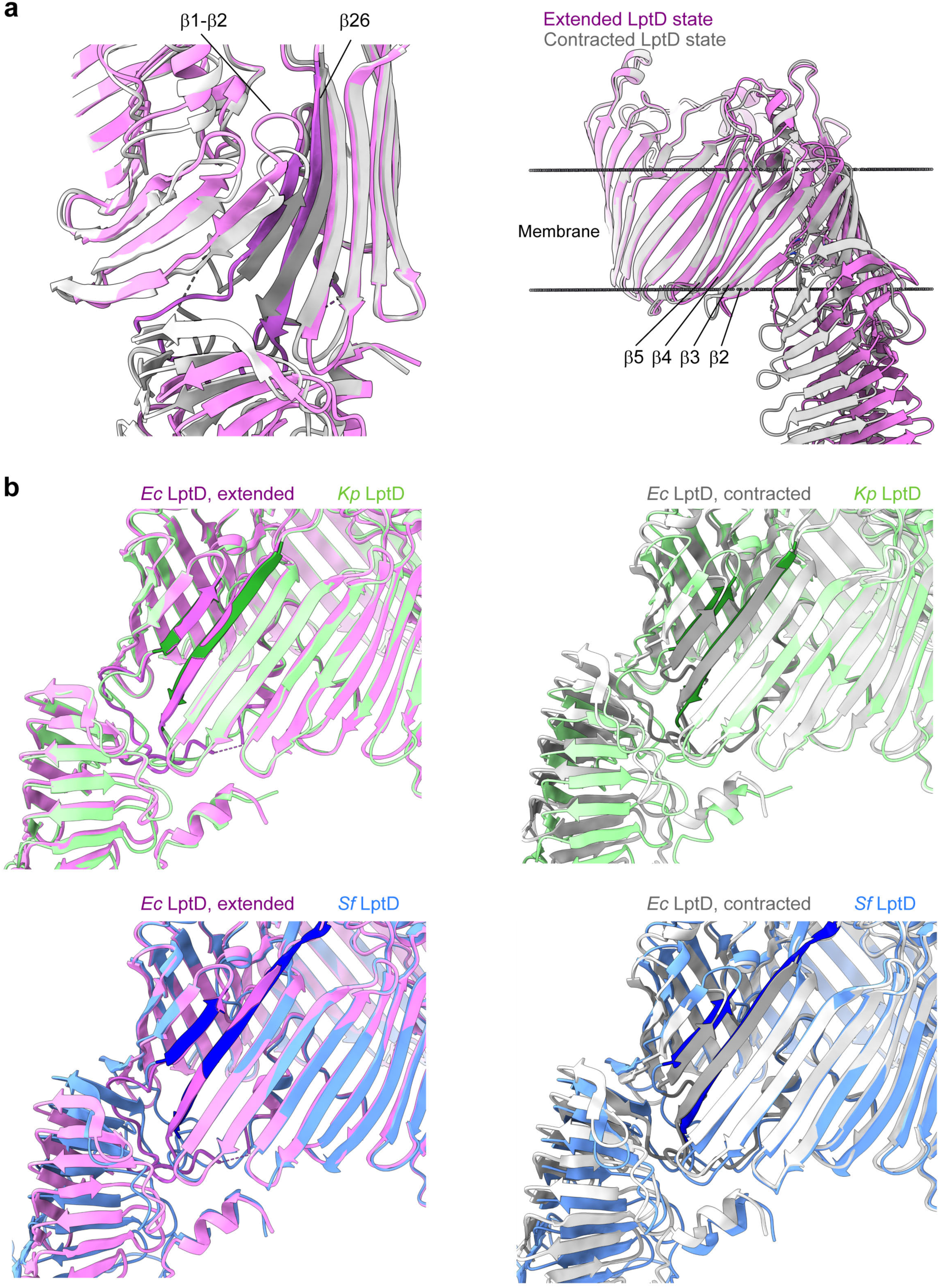
Differential features of LptD states. **a, Left,** Superimposition of the structure of LptD extended (purple) and contracted (grey). β1 and β26 are shown in darker colours. **Right,** Superimposed LptD extended (purple) and contracted (grey) are predicted for membrane topology using the OPM database server ^49^. The horizontal lines of grey dots represent the surfaces of a membrane lipid bilayer. **b**, Superimposition of the structure of LptD in the extended (purple) or contracted (grey) states with the structures of LptD from *Klebsiella pneumoniae* (green) or *Shigella flexneri* (blue). β1 and β26 are shown in darker colours.

**Extended Data Fig. 12.**
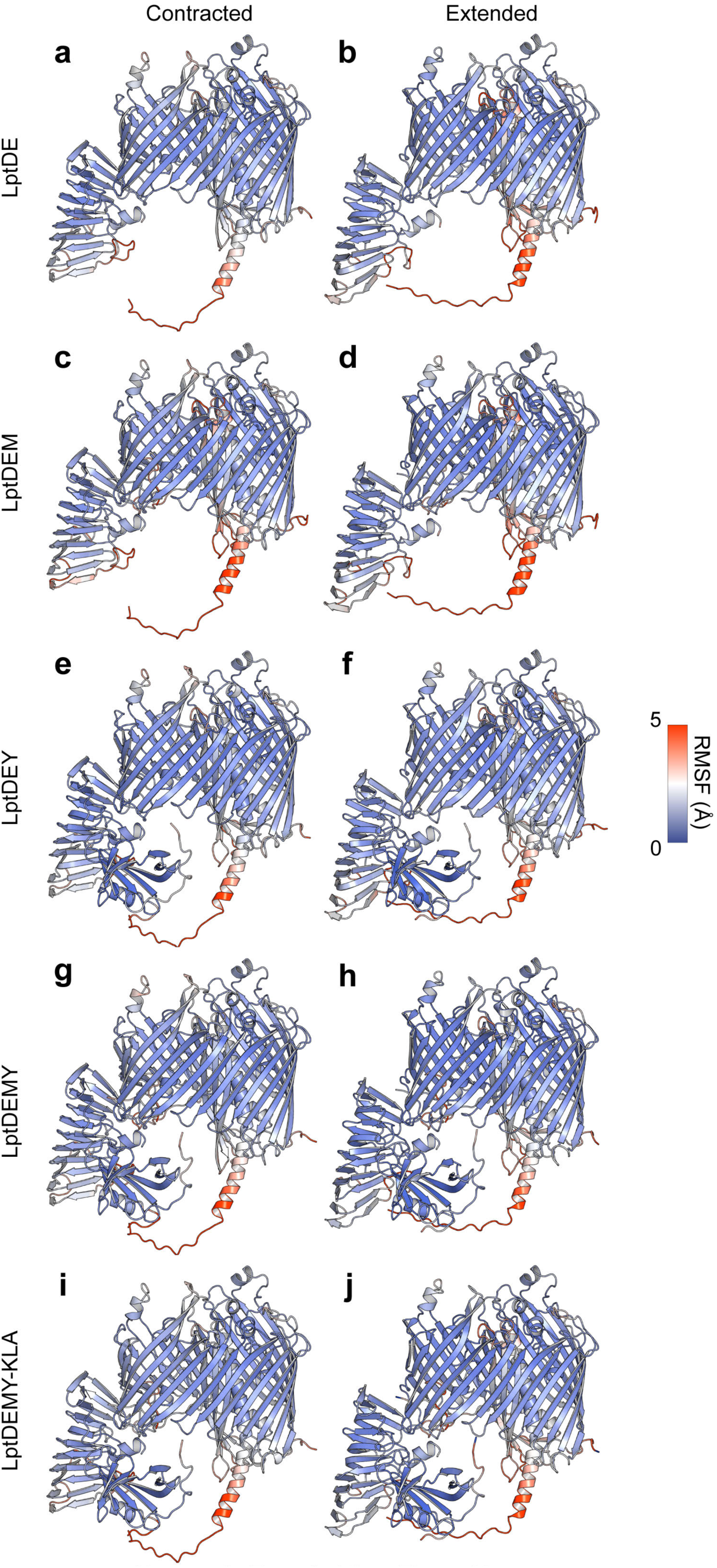
RMSF of simulated complexes. Cα root mean square fluctuations (RMSF) of the protein complexes simulated in this study. Data is shown on a blue to red scale and coloured onto a protein ribbon representation. Simulations are shown for the five contracted (**a**, **c**, **e**, **g**, and **i**) and extended (**b**, **d**, **h**, and **j**) states for (a, b) LptDE, (c, d) LptDEM, (e, f,) LptDEY, (g, h) LptDEMY, and (l, j) LptDEMY with bound KLA.

**Extended Data Fig. 13.**
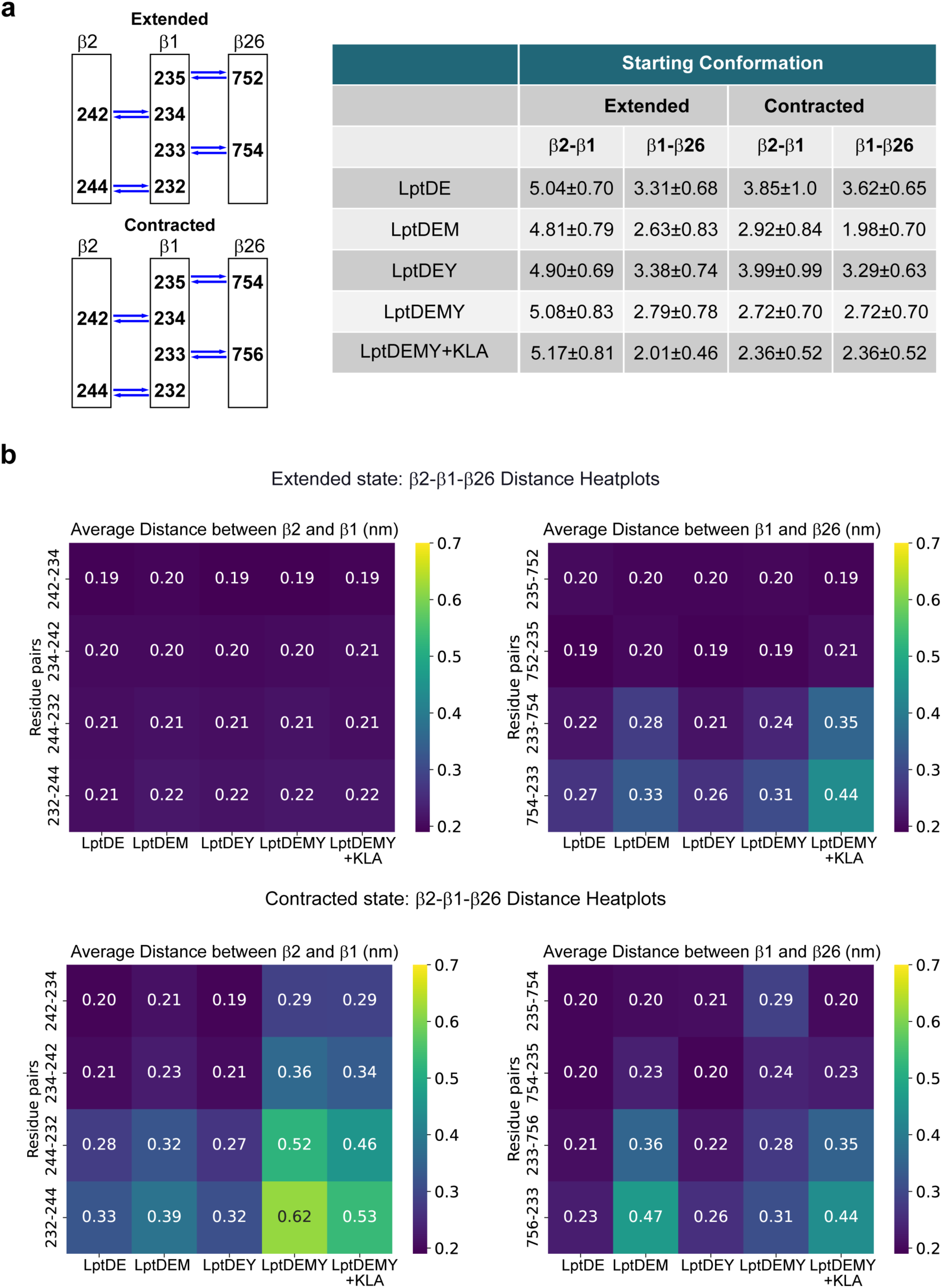
β2-β1-β26 pairing during MD simulations. **a**, **Left**, diagram illustrating β2-β1-β26 pairing in the extended and contracted LptD states. **Right**, average number of H-bonds computed during MD simulations for LptD in the extended state (LptDEM) or contracted state (LptDEY) upon addition or removal of LptM or LptY or addition of KLA as indicated in each row (See also Extended Data Figure 12). **b**, Average distances between β2-β1 or β1-β26 calculated for the indicated complexes during MD simulations described in in a).

**Extended Data Fig. 14.**
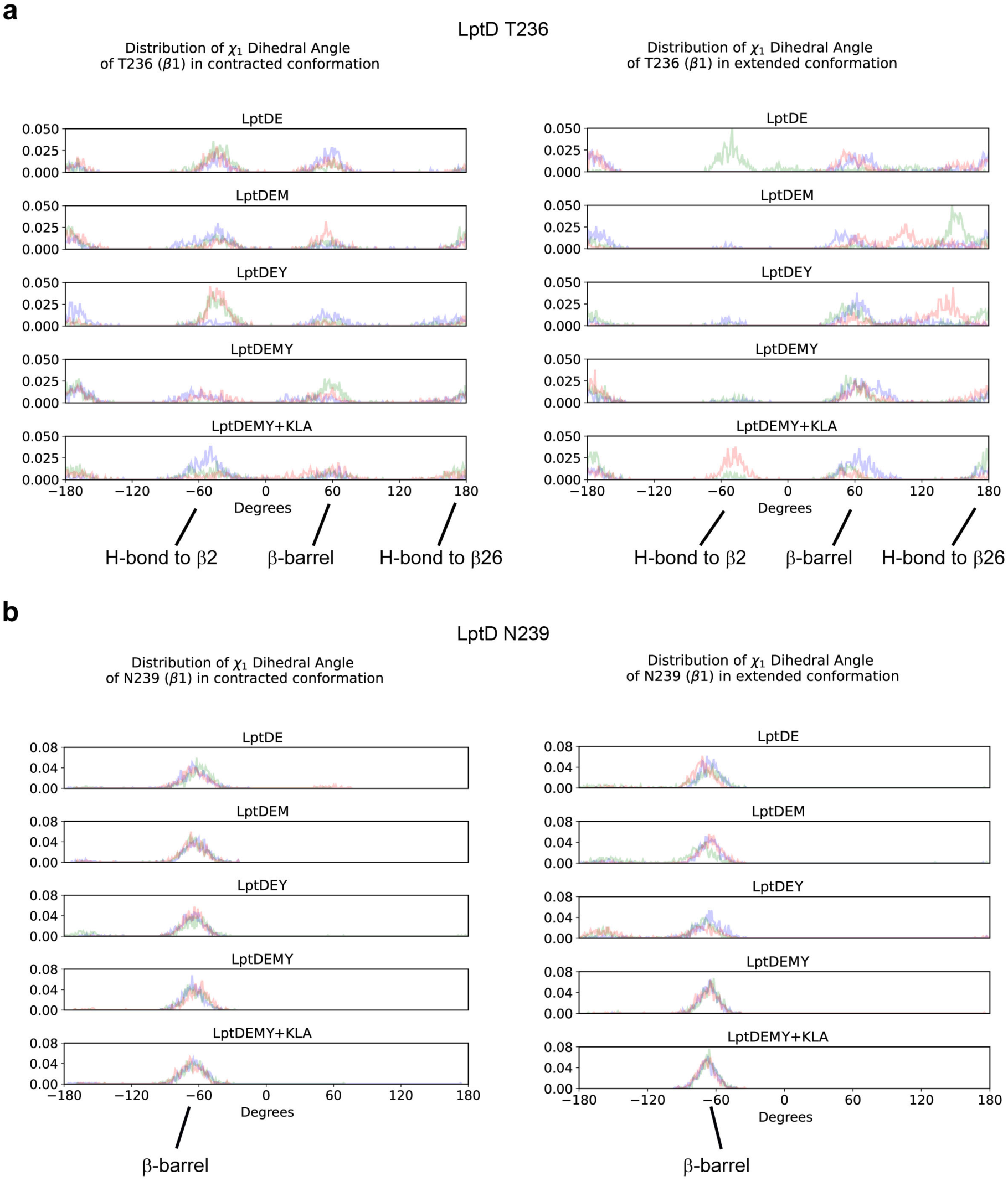
T236 and N239 dihedral angle distributions. Distribution of dihedral angle of LptD T236 (**a**) and LptD N239 (**b**) during three independent MD simulations runs.

**Extended Data Table 1.** LptD – LptM contacts. The interface contacts between LptD and LptM were determined in ChimeraX by identifying the pairs of atoms between the experimental model of LptD and the AlphaFold 2 in silico model of LptY with a Van der Waals overlap > −0.4 A.

| LptDEM |  |  |  |  | LptDEMY |  |  |  |  |
| --- | --- | --- | --- | --- | --- | --- | --- | --- | --- |
| LptM # | LptM AA | LptD # | LptD AA | contacts | LptM # | LptM AA | LptD # | LptD AA | contacts |
| 20 | CYS | 216 | LEU | 1 |  |  |  |  |  |
| 20 | CYS | 217 | GLN | 4 | 20 | CYS | 217 | GLN | 3 |
| 20 | CYS | 219 | PRO | 1 |  |  |  |  |  |
| 20 | CYS | 228 | PHE | 3 | 20 | CYS | 228 | PHE | 3 |
| 21 | GLY | 219 | PRO | 1 | 21 | GLY | 219 | PRO | 1 |
| 21 | GLY | 225 | ARG | 4 | 21 | GLY | 225 | ARG | 4 |
| 21 | GLY | 227 | GLY | 1 | 21 | GLY | 227 | GLY | 1 |
| 21 | GLY | 228 | PHE | 3 | 21 | GLY | 228 | PHE | 3 |
|  |  |  |  |  | 21 | GLY | 229 | LEU | 4 |
| 22 | LEU | 219 | PRO | 5 | 22 | LEU | 219 | PRO | 4 |
| 22 | LEU | 221 | GLY | 1 | 22 | LEU | 221 | GLY | 1 |
| 22 | LEU | 223 | LYS | 2 | 22 | LEU | 223 | LYS | 1 |
| 22 | LEU | 224 | ARG | 2 | 22 | LEU | 224 | ARG | 4 |
| 22 | LEU | 225 | ARG | 6 | 22 | LEU | 225 | ARG | 4 |
| 22 | LEU | 248 | TYR | 6 | 22 | LEU | 248 | TYR | 3 |
|  |  |  |  |  | 23 | LYS | 224 | ARG | 1 |
| 23 | LYS | 248 | TYR | 8 | 23 | LYS | 248 | TYR | 9 |
| 23 | LYS | 258 | THR | 2 | 23 | LYS | 258 | THR | 1 |
| 23 | LYS | 275 | GLU | 4 | 23 | LYS | 275 | GLU | 4 |
| 24 | GLY | 224 | ARG | 5 | 24 | GLY | 224 | ARG | 5 |
| 24 | GLY | 277 | ARG | 3 | 24 | GLY | 277 | ARG | 3 |
| 25 | PRO | 224 | ARG | 3 | 25 | PRO | 224 | ARG | 4 |
|  |  |  |  |  | 26 | LEU | 224 | ARG | 4 |
| 26 | LEU | 253 | PRO | 1 | 26 | LEU | 253 | PRO | 2 |
| 26 | LEU | 254 | ASN | 2 | 26 | LEU | 254 | ASN | 2 |
| 26 | LEU | 255 | MET | 3 | 26 | LEU | 255 | MET | 1 |
| 26 | LEU | 256 | ASP | 3 | 26 | LEU | 256 | ASP | 1 |
| 26 | LEU | 277 | ARG | 4 | 26 | LEU | 277 | ARG | 2 |
| 26 | LEU | 278 | TYR | 3 | 26 | LEU | 278 | TYR | 4 |
|  |  |  |  |  | 26 | LEU | 279 | LEU | 1 |
| 27 | TYR | 253 | PRO | 10 | 27 | TYR | 224 | ARG | 3 |
| 27 | TYR | 254 | ASN | 4 | 27 | TYR | 253 | PRO | 10 |
| 27 | TYR | 279 | LEU | 3 | 27 | TYR | 254 | ASN | 3 |
| 28 | PHE | 279 | LEU | 7 | 28 | PHE | 279 | LEU | 5 |
| 28 | PHE | 280 | SER | 2 | 28 | PHE | 280 | SER | 2 |
| 29 | PRO | 253 | PRO | 2 | 29 | PRO | 253 | PRO | 3 |
| 29 | PRO | 254 | ASN | 2 | 29 | PRO | 254 | ASN | 1 |
| total |  |  |  | 107 | total |  |  |  | 107 |

**Extended Data Table 2.** LptD – LptY contacts. The interface contacts between LptD and LptY were determined in ChimeraX by identifying the pairs of atoms between the experimental model of LptD and the AlphaFold 2 in silico model of LptY with a Van der Waals overlap > −0.4 A.

| LptDEY |  |  |  |  |
| --- | --- | --- | --- | --- |
| LptY # | LptY AA | LptD # | LptD AA | contacts |
| 16 | CYS | 646 | ARG | 27 |
| 17 | ALA | 646 | ARG | 1 |
| 18 | GLU | 619 | ARG | 14 |
| 18 | GLU | 646 | ARG | 3 |
| 48 | VAL | 30 | GLN | 4 |
| 48 | VAL | 174 | LEU | 4 |
| 68 | ARG | 33 | LEU | 8 |
| 69 | GLN | 33 | LEU | 1 |
| 70 | TRP | 33 | LEU | 4 |
| 70 | TRP | 34 | GLY | 8 |
| 70 | TRP | 174 | LEU | 6 |
| 70 | TRP | 175 | PRO | 19 |
| 71 | GLN | 33 | LEU | 2 |
| 71 | GLN | 34 | GLY | 13 |
| 71 | GLN | 36 | PRO | 1 |
| 71 | GLN | 175 | PRO | 4 |
| 72 | ARG | 32 | MET | 18 |
| 72 | ARG | 33 | LEU | 1 |
| 72 | ARG | 35 | VAL | 4 |
| 72 | ARG | 37 | SER | 5 |
| 74 | ILE | 29 | SER | 2 |
| 74 | ILE | 33 | LEU | 4 |
| 95 | GLU | 25 | ALA | 3 |
| 97 | TYR | 25 | ALA | 4 |
| total |  |  |  | 133 |

| LptDEMY |  |  |  |  |
| --- | --- | --- | --- | --- |
| LptY # | LptY AA | LptD # | LptD AA | contacts |
| 18 | GLU | 619 | ARG | 9 |
| 18 | GLU | 646 | ARG | 11 |
| 48 | VAL | 30 | GLN | 2 |
| 48 | VAL | 33 | LEU | 1 |
| 48 | VAL | 174 | LEU | 3 |
| 68 | ARG | 33 | LEU | 8 |
| 69 | GLN | 33 | LEU | 2 |
| 70 | TRP | 33 | LEU | 2 |
| 70 | TRP | 34 | GLY | 3 |
| 70 | TRP | 174 | LEU | 3 |
| 70 | TRP | 175 | PRO | 8 |
| 71 | GLN | 33 | LEU | 1 |
| 71 | GLN | 34 | GLY | 6 |
| 71 | GLN | 36 | PRO | 1 |
| 71 | GLN | 141 | GLN | 3 |
| 71 | GLN | 175 | PRO | 4 |
| 72 | ARG | 32 | MET | 5 |
| 72 | ARG | 35 | VAL | 6 |
| 72 | ARG | 36 | PRO | 1 |
| 72 | ARG | 37 | SER | 3 |
| 74 | ILE | 29 | SER | 1 |
| 74 | ILE | 32 | MET | 1 |
| 74 | ILE | 33 | LEU | 6 |
| 97 | TYR | 25 | ALA | 4 |
| 109 | ASP | 25 | ALA | 6 |
| total |  |  |  | 94 |

## SUPPLEMENTARY INFORMATION

## METHODS

### Bacterial strains and growth conditions

The *E. coli* strains used in this study are listed in Supplementary Table 1. All deletion strains were generated by transducing disrupted alleles into BW25113 ^1^ using P1 phage lysates of the corresponding deletion strains in the Keio collection ^2^. To obtain double deletion strains the Kanamycin-resistance (Kan^R^) cassette introduced with the first transduction event was excised as previously described ^3^. To delete chromosomal *lptD*, first BW25113 was transformed with a plasmid expressing LptD and LptE^His^ (pLptDE^His^) thus generating a *lptD* diploid strain. Then, the chromosomal *lptD* gene was replaced with a Kan^R^ cassette using bacteriophage lambda recombination. A P1 phage lysate of the derived strain (lptD::Kan^R^) was used to inactivate endogenous *lptD* by P1 transduction in *lptD* diploid strains harbouring a plasmid-borne copy of wild-type *lptD* or mutated *lptD*. The obtained transduced colonies were streaked once on lysogenic broth (LB) agar plates prior to LptD functional tests based on strain survival upon streaking on LB agar containing or lacking the oxidant 4,4-dipyridyl disulfide (4-DPS).

*E. coli* strains were generally cultured on LB liquid media or LB agar plates without antibiotics or supplemented with 100 μg/ml ampicillin or 50 μg/ml kanamycin. Serial dilution assays were conducted on either LB agar or MacConkey agar plates supplemented with 120 μg/ml vancomycin, or 50 μM Isopropyl β-D-1-thiogalactopyranoside (IPTG), as indicated in the figures.

### Plasmid construction

Plasmids and oligonucleotides are listed respectively in Supplementary Tables 2 and 3. Plasmids harbouring *lptY* alone (pLptY^His^) or in operon with *lptD*, *lptE* or *lptM* (pLptDEY^His^, pLptD^β-barrel^EY^His^) downstream of a P*_lac_* derivative promoter were generated by integrating *lptY* in pTrc99a or in previously described vectors (pLptDEM^His^, pLptD^β-barrel^EM^His3^) using a recombination-based, ligation-free cloning protocol. Plasmids encoding truncated forms of LptM were also generated by recombination-based cloning or by inverse PCR (see Supplementary Tables 2 and 3) using pLptDME^His^ as template. The replacement of specific *lptD* codons by an amber codon or by a cysteine-encoding codon was performed either by site-directed mutagenesis or by recombination-based cloning (see Supplementary Table 2).

### Cell fractionation and isolation of protein complexes

Total cell lysates were prepared from cells cultured to mid-exponential phase at 37°C. Where indicated, cells were supplemented with IPTG for 1.5h prior to sample collection by centrifugation, lyses in Laemmli buffer and boiling. Protein affinity purification under mild-solubilization conditions was conducted as previously described ^4^. Briefly, protein expression was induced in cell cultures at mid-exponential phase (OD600=0.5) by supplementing 200 μM IPTG for 1.5h prior to cell collection by centrifugation. Cells were resuspended in 20 mM Tris-HCl pH 8 containing an EDTA-free protease inhibitor cocktail (Roche). This buffer was further supplemented with 50mM iodoacetamide to alkylate free thiol groups in cysteine residues of proteins that had to be analysed for the presence of disulfide bonds. Resuspended cells were mechanically disrupted using a cell disruptor (Constant Systems LTD) set to 0.9 kPa. The obtained cell lysate was clarified by centrifugation at 6,000 x *g*, 4°C for 30 min. The crude envelope fraction was collected by subjecting the supernatant to ultracentrifugation at 50,000 x *g* at 4 °C for 30 min. The crude envelope fraction was solubilized with 50 mM Tris-HCl pH 8, 150mM NaCl, 20 mM imidazole supplemented with EDTA-free protease inhibitor (Roche) and 1% (w/v) n-dodecyl-β-D-maltopyranoside (DDM, Merck). After a clarifying spin to remove insoluble material, solubilized proteins were incubated with Protino Ni-NTA resin (Machery-Nagel) for 2h at 4°C. After extensive washes of the column with 50 mM Tris-HCl pH 8, 150mM NaCl, 50 mM imidazole and 0.03% (w/v) DDM, bound proteins were eluted with a similar buffer containing 800 mM imidazole. Aliquots of the elution fractions supplemented with 10% (w/v) glycerol were snap-frozen in liquid nitrogen for storage at −80 °C or directly analyzed by SDS or blue native gel electrophoresis. The purification of LptDEM^His^ and LptDE^His^ for cryo-EM complexes was performed from 2-liter cultures of wild-type and τι*lptM* strains carrying respectively pLptDEM^His^ or pLptDE^His^. Membranes were prepared and solubilized in DDM as describe above. Protein complexes were first purified by nickel-affinity chromatography and then subjected to size exclusion chromatography (Superdex 200 increase 10/300, Cytiva) in 50 mM Tris-HCl pH 8, 50mM NaCl and 0.03% (w/v) DDM. Fractions of interest were collected and concentrated to 0.3 mg/mL of proteins by ultrafiltration with a molecular weight cut-off of 100 kDa (Millipore). Fresh protein samples were immediately used to prepare cryo-EM grids.

### Site-directed photocrosslinking

Site directed photocrosslinking was performed as previously described ^3^ using wild type and τι*lptM* cells transformed with pEVOL-pBpF and variants of pLptDE^His^ or pLptDEM^His^ harbouring amber codons at specific positions in the reading frame of *lptD* (Supplementary Table 2). Transformed cells were cultured in minimal media until mid-exponential phase, supplemented with 1 mM pBpa and 200 μM IPTG for 1.5h. Two identical cell culture aliquots were withdrawn, one was kept on ice protected from light and the other was subjected to UV irradiation (Tritan 365 MHB, Spectroline) for 10 min on ice. Envelope fractions prepared from both non-treated and UV irradiated cells were subjected to DDM-solubilization and nickel-affinity purification of LptE^His^ or LptM^His^ to isolated LptDE or LptDEM complexes as described above.

### Gel electrophoresis and protein detection

Proteins samples were diluted in Laemmli buffer lacking β-mercaptoethanol (non-reducing conditions) or supplemented with β-mercaptoethanol (reducing conditions). Proteins were separated by home-made SDS polyacrylamide gels (10% acrylamide in Bis-Tris pH 6.4 buffer, subjected to electrophoresis using either MES or MOPS buffer). Where indicated, gels were stained with Coomassie Brilliant Blue R250. To perform Western blots, protein gels were blotted onto PVDF membranes and subjected to immunodetection using epitope-specific rabbit polyclonal antisera or with an anti-LPS mouse monoclonal antibody (WN1 222-5, Hycult Biotech). Immunodetection was revealed by using enhanced chemiluminescence. The signal intensity of protein bands was quantified using Image Lab software (BioRad). The rabbit polyclonal antiserum against LptD and against LptA were kind gifts of Drs. J.F. Collet (Louvain, Belgium) and A. Polissi (Milan, Italy), respectively.

### Native MS

Graphs presented in Extended Data Fig. 4a-b were obtained by re-analysing data previously deposited to the ProteomeXchange Consortium via the PRIDE partner repository ^5^ with the dataset identifier PXD041774. MS spectra presented in Extended Data Fig. 4c-d and showing the dissociated monomers were acquired with similar parameters as in ^3^, but with an extended m/z range of 1,500-3,000 Th. Raw data were acquired with MassLynx 4.1 (Waters, Manchester, UK) and analysed manually.

### In-gel tryptic digestion

LptDEM^His^ and LptDE^His^ complexes purified for cryo-EM experiments were loaded and separated on 4-12% polyacrylamide gradient gel (Merck) using MES buffer. Protein bands were revealed by Coomassie Brilliant Blue staining. Bands of interest were excised from the gel, cut into small pieces to perform in-gel digestion as previously described ^6^. After several washing steps to eliminate stain, the pieces of gel were dried under vacuum. Proteins were reduced with 10 mM DTT in 100 mM ammonium bicarbonate buffer for 35 min at 56 °C and then alkylated with 55 mM iodoacetamide for 30 min at room temperature in the dark. The gel fragments were dried under vacuum and swollen in a covering volume of modified trypsin (Promega, Madison, WI, USA) solution (12.5 ng/ml in 50 mM ammonium bicarbonate buffer) for 15 min in an ice bath followed by overnight incubation at 37 °C. The supernatant was pooled with two peptide extracts performed at 37°C for 15 min with shaking in 5% formic acid in 50% acetonitrile. The peptide mixture was dried down with the speed vacuum concentrator and re-suspended in 20 μL of 0.05 % formic acid in 2 % acetonitrile for nano-LC-MS/MS analysis.

### LC-MS/MS analysis

Samples were analysed using an Ultimate 3000 nanoRS system coupled to a Q-Exactive Plus mass spectrometer (Thermo Fisher Scientific, Bremen, Germany) operating in positive mode. 5 µL of each sample was loaded onto a C18-precolumn (300 µm inner diameter x 5 mm) at 20 µL/min in 2% ACN, 0.05% trifluoroacetic acid (TFA). After 5 min of desalting, the precolumn was switched online with the analytical C18 column (75 μm inner diameter × 50 cm, in-house packed with Reprosil C18) equilibrated in 95% solvent A (5% ACN, 0.2% FA) and 5% solvent B (80% ACN, 0.2% FA). Peptides were eluted by using a 5–25% gradient of solvent B for 40 min, then a 25–50% of solvent B for 20 min at a flow rate of 300 nL/min. The Q-Exactive Plus was operated in data-dependent acquisition mode. Survey scans MS were acquired in the Orbitrap over 300–2,000 m/z with a resolution of 70,000 (at m/z 400) an automatic gain control (AGC) target value of 3e6, and a maximum injection time of 100ms. The 10 most intense multiply charged ions (up to 2+) were selected at 2 m/z and fragmented by Higher Energy Collisional Dissociation (normalized collision energy set to 27). The resulting fragments were analysed in the Orbitrap with a resolution of 17,500 (at 400 m/z), an automatic gain control (AGC) target value of 1e5 and a maximum injection time of 50ms. Dynamic exclusion was used within 30 s with a 10 ppm tolerance, to prevent repetitive selection of the same peptide. For internal calibration, the 445.120025 ion was used as lock mass.

### Bioinformatic MS data analysis

Acquired MS and MS/MS data as raw MS files were converted to the mzDB format ^7^ using the pwiz-mzdb converter (version 0.9.10, https://github.com/mzdb/pwiz-mzdb) executed with its default parameters. Generated mzDB files were processed with the mzdb-access library (version 0.7, https://github.com/mzdb/mzdb-access) to generate peaklists. Peak lists were searched against SwissProt protein database with taxonomy *Escherichia coli* (23,135 sequences) in Mascot search engine (version 2.8.3, Matrix Science, London, UK). Cysteine carbamidomethylation was set as a fixed modification. Methionine oxidation and acetylation of protein N-terminus were set as variable modification. Up to two missed trypsin/P cleavages were allowed. Mass tolerances in MS and MS/MS were set to 10 ppm and 20 mmu, respectively. Proline software ^8^ was used for the validation and the label-free quantification of identified proteins in each sample. Mascot identification results were imported into Proline. Search results were validated with a peptide rank=1 and at 1 % FDR both at PSM level (on Adjusted e-Value criterion) and protein sets level (on Modified Mudpit score criterion). Label-free quantification was performed for all proteins identified: peptides were quantified by extraction of MS signals in the corresponding raw files, and post-processing steps were applied to filter, normalize, and compute protein intensities. The cross-assignment of MS/MS information between runs was enabled (it allows to assign peptide sequences to detected but non-identified features). Each protein intensity was based on the sum of unique peptide intensities and was normalized across all samples by the median intensity. Protein abundances were summarized in iBAQ values by dividing the protein intensities by the number of observable peptides in order to determine the protein stoichiometry ^9,10^.

### Cryo-EM sample preparation and data acquisition

2 µL of sample were deposited onto glow-discharged R2/1 carbon grids with an additional 2 nm ultrathin continuous carbon layer and placed in the thermostatic chamber of a Leica EM-GP automatic plunge freezer, set at 4°C and 95% humidity. Excess solution was removed by blotting with Whatman n°1 filter paper and the grids were immediately flash frozen in liquid ethane at −185°C. Images were pre-screened in house on a Talos Arctica (Thermo Fisher Scientific) operated at 200kV in parallel beam condition with a K3 Summit direct electron detector and a BioQuantum energy filter (Gatan Inc.). Energy-filtered (20 eV slit width) image series were acquired with Digital Micrograph software.

Data collection was performed at EMBL Heidelberg on a Titan Krios G4 microscope equipped with a cold field emission gun (C-FEG) operated at 300 kV. Images were recorded using a Falcon 4i direct electron detector in counting mode, coupled to a SelectrisX energy filter with a 10 eV energy slit width. SerialEM (version 4.1.0beta) ^11^ was used for automated data collection at a nominal magnification of 165,000x in nanoprobe mode (spot size 5). The microscope was operated with a 50 μm C2 aperture and a 100 µm objective aperture, with a beam diameter of 480 nm and a spherical aberration (Cs) of 2.7 mm. This resulted in a calibrated pixel size of 0.731 Å at the specimen level.

### Image processing

All image processing steps were performed in cryoSPARC v4 ^12^. EER movies from both datasets were imported and processed using Patch motion correction, while CTF parameters were estimated using Patch CTF estimation. After initial inspection, micrographs showing significant drift, poor CTF fits, or ice contamination were discarded.

### Determination of LptDE and LptDEY structures

Initial data preprocessing was performed on 20,832 micrographs in cryoSPARC. After CTF estimation and manual curation, 20,789 micrographs were selected for further processing. Initial particle picking was performed using a blob picker with a diameter range of 100-150 Å, yielding 7,573,562 particles, from which 5,280,207 particles were retained after inspection. A subset of 1,116,177 particles from 5,000 micrographs was extracted with a box size of 400 pixels for initial processing.

After 2D classification into 100 classes, 337,670 particles (30%) were selected and further classified into 50 classes, yielding 180,417 particles (53%). Based on these initial classes, a second round of template-based particle picking was performed using a 170 Å diameter, resulting in 11,445,246 particles. After inspection and cleanup, 5,219,710 particles were extracted. These particles underwent multiple rounds of 2D classification: first into 100 classes yielding 1,916,647 particles (37%), then into 50 classes resulting in 829,104 particles (43%), and finally to 229,126 particles (28%) showing high-quality features.

The selected particles were subjected to *ab initio* reconstruction followed by homogeneous refinement, reaching 4.25 Å resolution. After local refinement, the resolution improved to 2.89 Å. The particles were then re-extracted with a 360-pixel box size and underwent multiple rounds of heterogeneous refinement, progressively improving the particle set through 187,497, 144,168, and finally 86,702 particles.

A final round of template-based particle picking was performed with a 160 Å diameter. After extraction and extensive 2D classification, a set of 457,205 particles was obtained and refined to 3.18 Å, with local refinement improving the resolution to 2.44 Å. Ab initio reconstruction into four classes followed by further refinement produced the final LptDE reconstruction at 2.74 Å resolution from 107,410 particles. Resolution was estimated using the FSC=0.143 criterion.

A parallel processing branch starting with 829,104 particles underwent heterogeneous refinement yielding 361,218 particles, then 254,562 particles, and finally 187,497 particles. After additional refinement and 2D classification steps, 86,702 particles were obtained. A final round of template picking was performed using a 160 Å diameter. The resulting 13,173,675 particles underwent multiple rounds of processing, including extraction and binning (360 px → 180 px) of 6,375,528 particles, followed by successive 2D classifications with 200 and 100 classes. The final set of 127,785 particles was refined to 2.94 Å, with local refinement yielding the final LptDEY reconstruction at 2.62 Å resolution. Resolution was estimated using the FSC=0.143 criterion.

### Determination of LptDEM and LptDEMY structures

Initial data preprocessing of 18,816 micrographs was performed in cryoSPARC. After manual curation, 18,287 micrographs (97%) were selected for further processing. Template-based particle picking was performed using a diameter of 250 Å, yielding 2,406,295 particles after inspection and extraction with a box size of 360 pixels. The extracted particles underwent two rounds of 2D classification. The first round with 100 classes yielded 909,602 particles (38%), which were further classified into 50 classes, resulting in a selection of 510,229 particles (56%). These particles were subjected to heterogeneous refinement with two volumes, and one population of 273,204 particles was refined to 2.86 Å using non-uniform refinement. After further classification and refinement steps, a set of 302,650 particles underwent three rounds of heterogeneous refinement, progressively improving the quality of the reconstruction. The final population of 198,109 particles was refined using homogeneous refinement to 2.63 Å, followed by local refinement that yielded the final LptDEM reconstruction at 2.47 Å resolution. Resolution was estimated using the FSC=0.143 criterion. For LptDEMY, further optimization focused on the dataset containing 302,650 particles. It underwent 3D classification into two classes using a mask focusing on the known location of LptY in the LptDEY structure. The particles belonging to the class displaying a density at the location of LptY (146,385 particles) were selected for final non-uniform refinement, yielding a reconstruction at 2.63 Å resolution. Resolution was estimated using the FSC=0.143 criterion.

### Model building and refinement

Initial models for LptD and LptE were generated using AlphaFold2 predictions ^13^ and rigid-body fitted into the experimental density maps. These initial models were modified through multiple rounds of manual rebuilding in ISOLDE ^14^ and COOT ^15^ to accurately fit the experimental density. For model building, maps were sharpened using DeepEMhancer ^16^ to enhance interpretability of structural features. LptM was built de novo into the corresponding density. For LptY-containing complexes, an AlphaFold2-predicted model was rigid-body docked into the experimental density. For real-space refinement in PHENIX ^17^, maps were sharpened using phenix.autosharpen. The models underwent multiple rounds of manual refinement in ISOLDE and real-space refinement using phenix.real_space_refine with appropriate geometric restraints. Final models were validated using MolProbity ^18^ and phenix.validation_cryoem implemented in the PHENIX software. Details about the cryo-EM refinement statistics and FSC Map versus Model plot can be found in Supplementary Table 4 and Extended Data Fig. 6, respectively

### Molecular Modelling and Dynamics simulations

LptM was added to an extended complex of LptDEY through superimposition of the extended and contracted states. The aligned coordinates of LptM were taken from the extended conformation and added to that of the contracted state. The final complex was then subjected to energy minimisation using GROMACS ^19^. Complexes of LptDE, LptDEM, LptDEY and LptDEMY in both contracted and extended states were assembled into model outer membranes using our MemProtMD protocol ^20^. In brief, the systems were converted to a Coase-Grained (CG) Martini 3 force field representation using Martinize2 ^21,22^. Here an elastic network with a force constant of 1000 kJ mol^−1^ nm^−2^ was used to connect Cα atoms within 10 Å in the protein structure. The protein was placed into a model outer membrane using *insane*, with 100% Kdo2-lipid A (KLA) in the outer leaflet and a 70:20:10 ratio of 1-palmitoyl-2-oleoyl-sn-glycero-3-phosphoethanolamine (POPE), 1-palmitoyl-2-oleoyl-sn-glycero-3-phosphoglycerol (POPG) and cardiolipin in the inner leaflet. The system was then solvated and ionized with 0.15M NaCl. The system was equilibrated in CG before conversion to a CHARMM36m ^23^ representation using CG2AT ^24^. Three repeats of 500 ns atomistic MD simulations were performed with a timestep of 2 fs for each LptDE assembly and state. All simulations were performed in the isothermal-isobaric ensemble at 310 K and 1 bar. Pressure was maintained at 1 bar with an semi-isotropic compressibility 3 x 10^−4^ using the C-rescale barostat^25^. Temperature was controlled using the velocity-rescale thermostat^26^, with the solvent, lipids and protein coupled to an external bath. All MD simulations were performed using GROMACS 2023 ^19^ and analysed using GROMACS tools and MDAnalysis ^27^. All images were generated using PyMOL ^28^. MD simulations data are available for all Lpt states and complexes for start (0ns) and end (500ns) frames for each simulation repeat (Zenodo repository, doi: 10.5281/zenodo.14834225). Starting tpr files are also available, and xtc trajectories can be supplied on request

### Genome samples

A representative set of *Enterobacterales* genomes was assembled by selecting genomes at genus (157 genomes) and species level (848 genomes) from the Genome Taxonomy Database (GTDB release 214) ^29^, which were downloaded from the NCBI website (ftp://ftp.ncbi.nlm.nih.gov/genomes). Genomes were annotated with *Prokka* in fast mode with default settings ^30^.

### Orthologous gene clusters and species tree

*OrthoFinder* software ^31^ was used to identify orthologous gene clusters (OGs) in the 157 representative genomes *of the Enterobacterales* genera. Before inferring the species tree, the 13 genomes with the fastest rates of evolution were discarded. 277 OGs, with zero or one gene copy per genome, were selected. Sequences were aligned with *mafft* (option -M msa) ^32^. Alignment sites with a gap frequency greater than 50% were removed and alignments concatenated. These steps were performed by OrthoFinder. The alignment contains 145 concatenated sequences and 80417 sites.

The tree was inferred with *IQ-TREE2* ^33^ and branch supports were assessed with ultrafast bootstrap approximation and with single branch tests (*-m LG+R10 -B 1000 - bnni -alrt 1000*). The tree was rooted with *Budviciaceae* as an outgroup and displayed with *iTOL* ^34^. The *Enterobacterales* family has been divided into seven clades: the *Enterobacteriaceae* clade, and the new families *Erwiniaceae*, *Pectobacteriaceae*, *Yersiniaceae*, *Hafniaceae*, *Morganellaceae* and *Budviciaceae* ^35^. In accordance with the species reported in this study, we generated a tree of the species present in our dataset. The branches, especially the deeper ones, are very well supported by the bootstrap values (Extended Data Fig. 2). The weakest branches are found in a subtree of the *Pectobacteriaceae*, whose leaves present very small genomes. The corresponding genera have no family assigned in the List of Prokaryotic names with Standing in Nomenclature (LPSN) ^36^ (*Schneideria*, *Baumannia*, *Mikella*, *Gullanella*, *Steffania*, *Hoaglandella*, and *Doolittlea*) or belong to *Enterobacteriaceae* (*Blochmannia* and *Moranella*). We used NCBI’s BioSample database to obtain genomes metadata.

### Search of proteins candidates

We chose LptDEM proteins as a signature for the LPS transporter in the *Enterobacterales* genomes. Annotation of the protein families were performed with *hmmscan* (*HMMER* package version 3.1b2) ^37^ and the Pfam library (version 34.0). Selected proteins were scanned against the Pfam library using *hmmscan* to discard false positives. Results were filtered to retain only alignments covering at least 75% of the Pfam domain and an independent E-value (i-Evalue) < 1e-15 for all but LptE and LptM (i-Evalue < 1e-5). Only one occurrence was retained per profile and per genome.

LptD is covered by two Pfam profiles (PF04453 and PF03968). The profile PF03968 is shared with LptA. Using a threshold of 50% for the percentage of profile coverage on the protein, we were able to distinguish between LptA and LptD proteins.

As previously observed ^3^, genes encoding the short lipoprotein LptM were not always annotated by *Prokka*. To complete the annotation, genomes for which no LptM had been identified were translated in all six reading phases with *esl-translate* program from *HMMER* package. *hmmsearch* was used to identify the presence of the PF13627 domain in translated ORFs (trORF). As above, *hmmscan* was used against the retained trORFs to remove false positives.

*E. coli* (strain K12) LptY (YEDD_ECOLI) is a lipoprotein of 137 amino acids long. LptY is characterized by Pfam PF13987. 359 and 58 sequences are selected from species and genus genome samples, respectively. We used the NCBI web server to search for YEDD_ECOLI in non-redundant protein sequences, excluding *Enterobacterales*. Both BlastP and BlastX software were used. The results show a small number of sequences (43) with high similarity to *E. coli* LptY. We aligned these sequences with those obtained from the *Enterobacterales* in our sample and inferred a tree. The non-*Enterobacterales* LptY proteins are identical or very close to *Enterobacterales* sequences and are generally partial. These observations indicate that these sequences are the result of either recent horizontal transfer or, more likely, sequencing artifacts.

### LptY tree

The 58 reference LptY protein sequences at genus level were aligned with *mafft* (--localpair --maxiterate 1000) ^32^. The filtered alignment with *trimAl* (-gappyout) included 136 amino acid sites ^38^. The LG4M substitution model was selected using *ModelFinder* ^39^. The tree was inferred with *IQ-TREE2* ^33^ and branch supports were estimated with ultra-fast bootstrap approximation, single branch tests (*-B 1000 -bnni - alrt 1000*) and standard non-parametric bootstrap (*-b 100*).

The genomic context of *lptY* gene candidates was extracted from GFF3 Prokka annotation files over a window 5000 nucleotides upstream and 10000 nucleotides downstream of these genes. The genes are associated with their HOG to illustrate the conservation of *lptY*’s genetic context. Pfam profiles associated with these HOGs were identified with *hmmscan* as described above.

### Analysis of *Enterobacterales* species tree

The tree is consistent with the new classification of the *Enterobacterales* family into seven clades ^35^: *Enterobacteriaceae*, *Erwiniaceae*, *Pectobacteriaceae*, *Yersiniaceae*, *Hafniaceae*, *Morganellaceae* and *Budviciaceae*; the branches, especially the deeper ones, are very well supported by the bootstrap values (Extended Data Fig. 2). The tree has been generated with genomes representing genera, but each genus may contain several genomes in our dataset (Extended Data Fig. 2, grey histogram). Intra-genus variability is illustrated by the size of the symbol, which is proportional to the frequency with which the feature is observed in the genomes of that genus. Genomes with significant genome reduction (Extended Data Fig. 2, genome size column) have accelerated sequence evolution, resulting in long branches. They almost always have arthropods as hosts, often in symbiotic associations (Extended Data Fig. 2, Metadata). The *Enterobacteriaceae* family includes almost all vertebrate pathogens but with low frequencies (small blue dots), which may reflect the poor metadata annotation of these genomes. The LptY protein has been identified in the *Enterobacteriaceae* and *Erwiniaceae* families, and with less frequency in the *Pectobacteriaceae* family. In *Erwiniaceae*, its absence is accompanied by the absence of other protein partners. In the Pectobacteriaceae, it is absent from the outer group, whose members often have Chromadorea (nematodes) as hosts. It is present in the first part of the other subtree, whose hosts are arthropods. There is a loss of LptM and, to a lesser extent, of LptD and LptE.

**Supplementary Table 1.** Bacterial strains.

| Strain name | Reference | Lab Identifier |
| --- | --- | --- |
| BW25113: $\Delta(araD-araB)567 \Delta(rhaD-$ | <sup>1</sup> | Y1 |
| *rhaB*)568 ΔlacZ4787(::rrnB-3) *hsdR*514 *rph*-1 (wild-type reference) | | |
| BW25113 *lptM*::kan | <sup>3</sup> | Y14 |
| BW25113 Δ*lptM* | <sup>4</sup> | Y40 |
| BW25113 *lptY*::kan | This study | Y-CHX51 |
| BW25113 Δ*lptY* | This study | Y-CHX15 |
| BW25113 Δ*lptM lptY*::kan | This study | Y-CHX49 |
| BW25113 *dsbA*::kan | <sup>3</sup> | Y81 |
| BW25113 Δ*lptY dsbA*::kan | This study | Y-CHX52 |
| BW25113 *lptD*::kan ; *pLptDE*<sup>His</sup> | This study | Y-P337 |

**Supplementary Table 2.** Plasmids.

| Plasmid name | Cloning strategies used in this study | Reference | Lab Identifier |
| --- | --- | --- | --- |
| pCtrl (empty vector) |  | <sup>4</sup> | pV3 |
| pLptDME<sup>His</sup> |  | <sup>3</sup> | pYY40 |
| pLptDM<sup>Δ22-30</sup>E<sup>His</sup> | Recombinational cloning | This study | pVM113 |
| pLptDM<sup>Δ44-67</sup>E<sup>His</sup> | Inverse PCR | This study | pVM90 |
| pLptDE<sup>His</sup> |  | <sup>3</sup> | pYY68 |
| pLptD<sup>K234C-N755C</sup>E<sup>His</sup> | Site-directed mutagenesis | This study | pVM186 |
| pLptD<sup>A233C-I756C</sup>E<sup>His</sup> | Site-directed mutagenesis | This study | pVM135 |
| pLptD<sup>G227C-Y248C</sup>E<sup>His</sup> | Site-directed mutagenesis | This study | pVM132 |
| pLptD<sup>Y112am</sup>E<sup>His</sup> | Site-directed mutagenesis | This study | pVM123 |
| pLptD<sup>F203am</sup>E<sup>His</sup> | Recombinational cloning | This study | pVM174 |
| pLptD<sup>V220am</sup>E<sup>His</sup> | Site-directed mutagenesis | This study | pVM124 |
| pLptD<sup>F228am</sup>E<sup>His</sup> | Site-directed mutagenesis | This study | pVM125 |
| pLptD<sup>I230am</sup>E<sup>His</sup> | Recombinational cloning | This study | pVM175 |
| pLptD<sup>N232am</sup>E<sup>His</sup> | Recombinational cloning | This study | pVM176 |
| pLptD<sup>T236am</sup>E<sup>His</sup> | Site-directed mutagenesis | This study | pVM126 |
| pLptD<sup>N239am</sup>E<sup>His</sup> | Site-directed mutagenesis | This study | pCHX33 |
| pLptDEM<sup>His</sup> |  | <sup>3</sup> | pYY43 |
| pLptD<sup>Y112am</sup>EM<sup>His</sup> | Site-directed mutagenesis | This study | pHX10 |
| pLptD<sup>F228am</sup>EM<sup>His</sup> | Site-directed mutagenesis | This study | pHX11 |
| pLptD<sup>T236am</sup>EM<sup>His</sup> | Site-directed mutagenesis | This study | pHX12 |
| pLptD<sup>β-barrel</sup>EM<sup>His</sup> |  | <sup>3</sup> | pVM52 |
| pLptD<sup>β-barrel</sup>E-LptY<sup>His</sup> | Recombinational cloning | This study | pCHX09 |
| pLptDE-LptY<sup>His</sup> | Recombinational cloning | This study | pCHX02 |
| pLptY<sup>His</sup> | Recombinational cloning | This study | pCHX01 |
| pDsbC |  | <sup>3</sup> | pVM41 |
| pLptC<sup>His</sup> |  | <sup>3</sup> | pVM83 |
| pEVOL-pBpF |  | <sup>40</sup> | pEVOL-pBpF |
| pKOBEG |  | <sup>41</sup> | pKOBEG |

**Supplementary Table 3.**
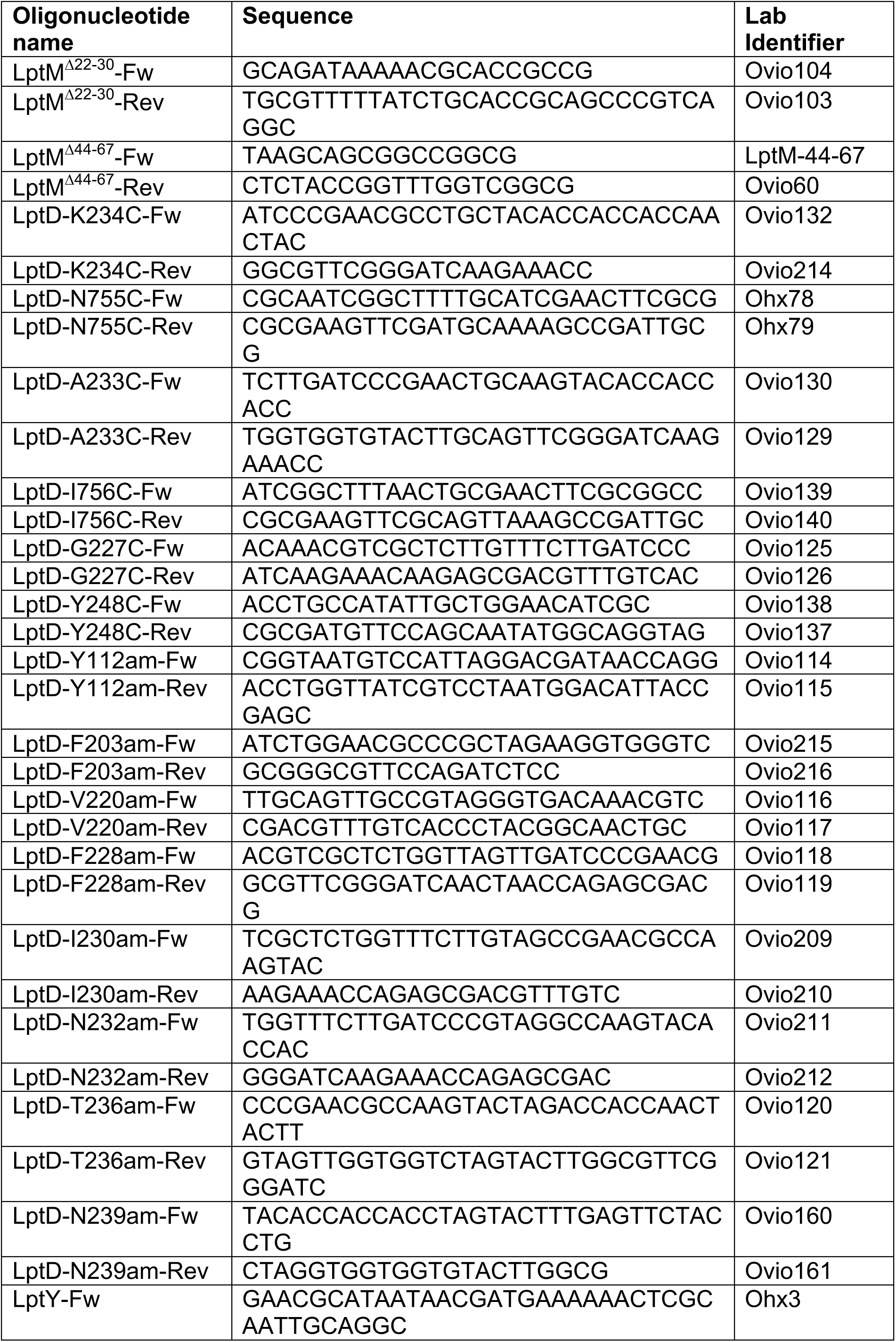
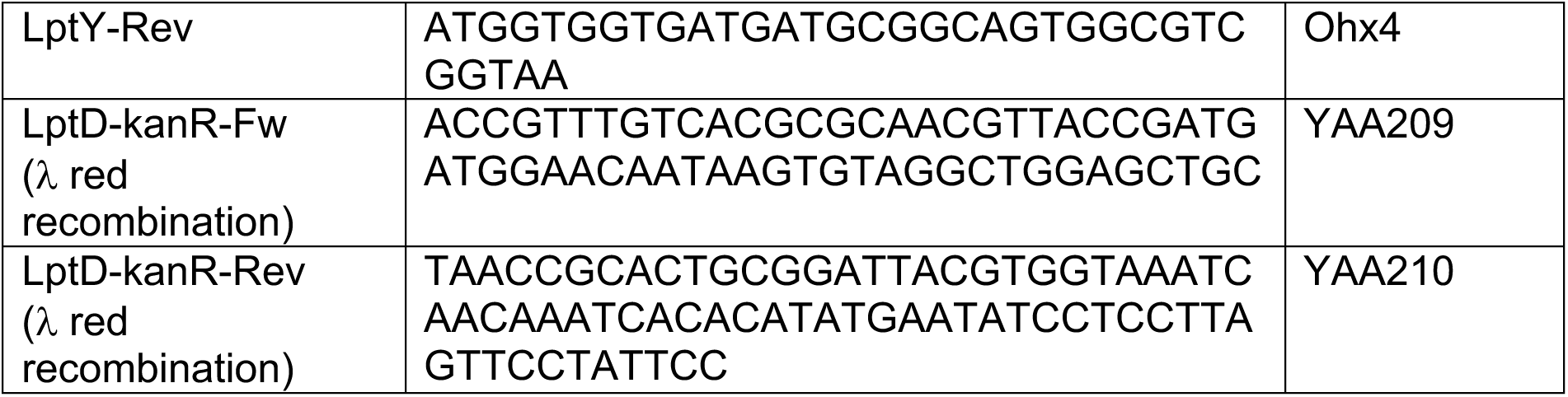
Oligonucleotides.

**Supplementary Table 4.**
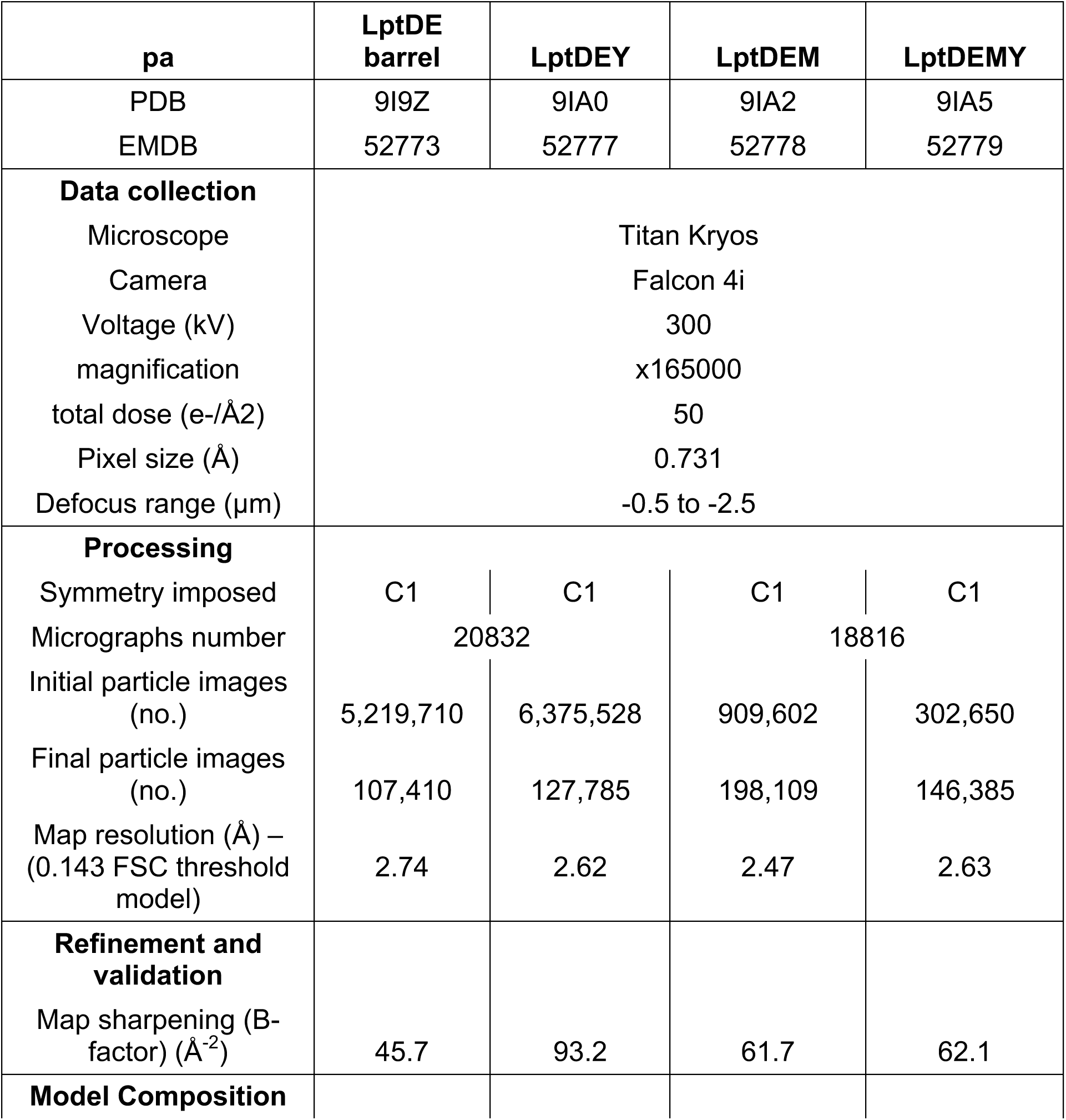
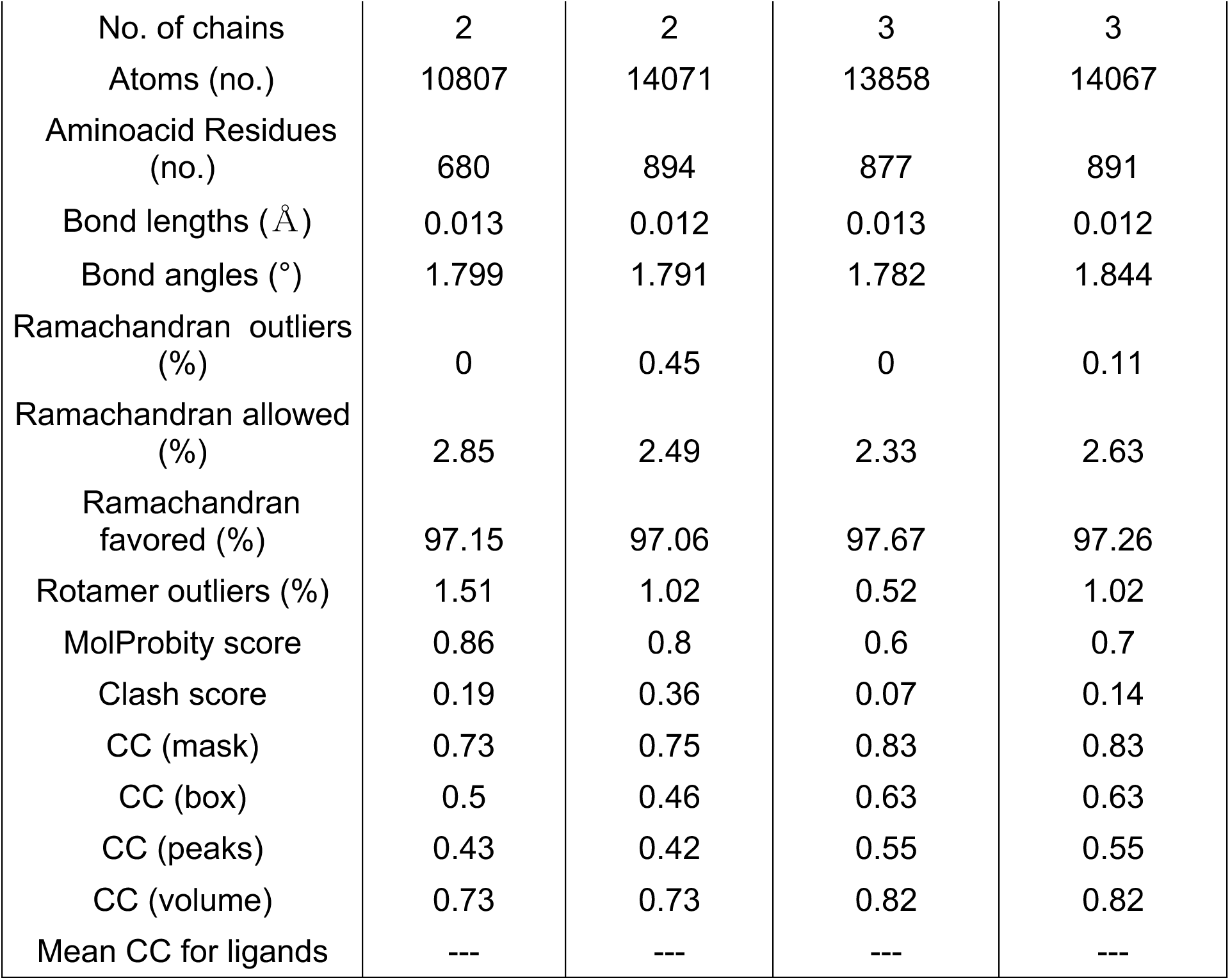
CryoEM data collection, processing and modelling statistics.

## REFERENCES

1 Lithgow, T., Stubenrauch, C. J. & Stumpf, M. P. H. Surveying membrane landscapes: a new look at the bacterial cell surface. Nature reviews. Microbiology 21, 502–518, doi:10.1038/s41579-023-00862-w (2023).

2 Sun, J., Rutherford, S. T., Silhavy, T. J. et al. Physical properties of the bacterial outer membrane. Nature reviews. Microbiology 20, 236–248, doi:10.1038/s41579-021-00638-0 (2022).

3 Hayashi, S., Buchanan, S. K. & Botos, I. The Name Is Barrel, β-Barrel. Methods in molecular biology (Clifton, N.J.) 2778, 1–30, doi:10.1007/978-1-0716-3734-0_1 (2024).

4 Witwinowski, J., Sartori-Rupp, A., Taib, N. et al. An ancient divide in outer membrane tethering systems in bacteria suggests a mechanism for the diderm-to-monoderm transition. Nature microbiology 7, 411–422, doi:10.1038/s41564-022-01066-3 (2022).

5 Heinz, E., Selkrig, J., Belousoff, M. J. et al. Evolution of the Translocation and Assembly Module (TAM). Genome biology and evolution 7, 1628–1643, doi:10.1093/gbe/evv097 (2015).

6 Doyle, M. T. & Bernstein, H. D. Molecular Machines that Facilitate Bacterial Outer Membrane Protein Biogenesis. Annual review of biochemistry 93, 211–231, doi:10.1146/annurev-biochem-030122-033754 (2024).

7 Sperandeo, P., Lau, F. K., Carpentieri, A. et al. Functional analysis of the protein machinery required for transport of lipopolysaccharide to the outer membrane of Escherichia coli. Journal of bacteriology 190, 4460–4469, doi:10.1128/jb.00270-08 (2008).

8 Wu, T., McCandlish, A. C., Gronenberg, L. S. et al. Identification of a protein complex that assembles lipopolysaccharide in the outer membrane of Escherichia coli. Proceedings of the National Academy of Sciences of the United States of America 103, 11754–11759, doi:10.1073/pnas.0604744103 (2006).

9 Bos, M. P., Tefsen, B., Geurtsen, J. et al. Identification of an outer membrane protein required for the transport of lipopolysaccharide to the bacterial cell surface. Proceedings of the National Academy of Sciences of the United States of America 101, 9417–9422, doi:10.1073/pnas.0402340101 (2004).

10 Sampson, B. A., Misra, R. & Benson, S. A. Identification and characterization of a new gene of Escherichia coli K-12 involved in outer membrane permeability. Genetics 122, 491–501, doi:10.1093/genetics/122.3.491 (1989).

11 Lundstedt, E., Kahne, D. & Ruiz, N. Assembly and Maintenance of Lipids at the Bacterial Outer Membrane. Chemical reviews 121, 5098–5123, doi:10.1021/acs.chemrev.0c00587 (2021).

12 Tan, W. B. & Chng, S. S. How Bacteria Establish and Maintain Outer Membrane Lipid Asymmetry. Annual review of microbiology 78, 553–573, doi:10.1146/annurev-micro-032521-014507 (2024).

13 Henderson, J. C., Zimmerman, S. M., Crofts, A. A. et al. The Power of Asymmetry: Architecture and Assembly of the Gram-Negative Outer Membrane Lipid Bilayer. Annual review of microbiology 70, 255–278, doi:10.1146/annurev-micro-102215-095308 (2016).

14 Rojas, E. R., Billings, G., Odermatt, P. D. et al. The outer membrane is an essential load-bearing element in Gram-negative bacteria. Nature 559, 617–621, doi:10.1038/s41586-018-0344-3 (2018).

15 Benn, G., Mikheyeva, I. V., Inns, P. G. et al. Phase separation in the outer membrane of Escherichia coli. Proceedings of the National Academy of Sciences of the United States of America 118, doi:10.1073/pnas.2112237118 (2021).

16 Ashraf, K. U., Nygaard, R., Vickery, O. N. et al. Structural basis of lipopolysaccharide maturation by the O-antigen ligase. Nature 604, 371–376, doi:10.1038/s41586-022-04555-x (2022).

17 Bertani, B. & Ruiz, N. Function and Biogenesis of Lipopolysaccharides. EcoSal Plus 8, doi:10.1128/ecosalplus.ESP-0001-2018 (2018).

18 Whitfield, C. & Trent, M. S. Biosynthesis and export of bacterial lipopolysaccharides. Annual review of biochemistry 83, 99–128, doi:10.1146/annurev-biochem-060713-035600 (2014).

19 Raetz, C. R. & Whitfield, C. Lipopolysaccharide endotoxins. Annual review of biochemistry 71, 635–700, doi:10.1146/annurev.biochem.71.110601.135414 (2002).

20 Okuda, S., Sherman, D. J., Silhavy, T. J. et al. Lipopolysaccharide transport and assembly at the outer membrane: the PEZ model. Nature reviews. Microbiology 14, 337–345, doi:10.1038/nrmicro.2016.25 (2016).

21 Laguri, C., Sperandeo, P., Pounot, K. et al. Interaction of lipopolysaccharides at intermolecular sites of the periplasmic Lpt transport assembly. Scientific reports 7, 9715, doi:10.1038/s41598-017-10136-0 (2017).

22 Törk, L., Moffatt, C. B., Bernhardt, T. G. et al. Single-molecule dynamics show a transient lipopolysaccharide transport bridge. Nature 623, 814–819, doi:10.1038/s41586-023-06709-x (2023).

23 Sherman, D. J., Xie, R., Taylor, R. J. et al. Lipopolysaccharide is transported to the cell surface by a membrane-to-membrane protein bridge. Science (New York, N.Y.) 359, 798–801, doi:10.1126/science.aar1886 (2018).

24 Botos, I., Majdalani, N., Mayclin, S. J. et al. Structural and Functional Characterization of the LPS Transporter LptDE from Gram-Negative Pathogens. Structure (London, England : 1993) 24, 965–976, doi:10.1016/j.str.2016.03.026 (2016).

25 Dong, H., Xiang, Q., Gu, Y. et al. Structural basis for outer membrane lipopolysaccharide insertion. Nature 511, 52–56, doi:10.1038/nature13464 (2014).

26 Qiao, S., Luo, Q., Zhao, Y. et al. Structural basis for lipopolysaccharide insertion in the bacterial outer membrane. Nature 511, 108–111, doi:10.1038/nature13484 (2014).

27 Freinkman, E., Chng, S. S. & Kahne, D. The complex that inserts lipopolysaccharide into the bacterial outer membrane forms a two-protein plug-and-barrel. Proceedings of the National Academy of Sciences of the United States of America 108, 2486–2491, doi:10.1073/pnas.1015617108 (2011).

28 Lee, J., Xue, M., Wzorek, J. S. et al. Characterization of a stalled complex on the β-barrel assembly machine. Proceedings of the National Academy of Sciences of the United States of America 113, 8717–8722, doi:10.1073/pnas.1604100113 (2016).

29 Ruiz, N., Chng, S. S., Hiniker, A. et al. Nonconsecutive disulfide bond formation in an essential integral outer membrane protein. Proceedings of the National Academy of Sciences of the United States of America 107, 12245–12250, doi:10.1073/pnas.1007319107 (2010).

30 Chng, S. S., Xue, M., Garner, R. A. et al. Disulfide rearrangement triggered by translocon assembly controls lipopolysaccharide export. Science (New York, N.Y.) 337, 1665–1668, doi:10.1126/science.1227215 (2012).

31 Denoncin, K., Vertommen, D., Paek, E. et al. The protein-disulfide isomerase DsbC cooperates with SurA and DsbA in the assembly of the essential β-barrel protein LptD. The Journal of biological chemistry 285, 29425–29433, doi:10.1074/jbc.M110.119321 (2010).

32 Gu, Y., Stansfeld, P. J., Zeng, Y. et al. Lipopolysaccharide is inserted into the outer membrane through an intramembrane hole, a lumen gate, and the lateral opening of LptD. Structure (London, England : 1993) 23, 496–504, doi:10.1016/j.str.2015.01.001 (2015).

33 Li, X., Gu, Y., Dong, H. et al. Trapped lipopolysaccharide and LptD intermediates reveal lipopolysaccharide translocation steps across the Escherichia coli outer membrane. Scientific reports 5, 11883, doi:10.1038/srep11883 (2015).

34 Lundquist, K. P. & Gumbart, J. C. Presence of substrate aids lateral gate separation in LptD. Biochimica et biophysica acta. Biomembranes 1862, 183025, doi:10.1016/j.bbamem.2019.07.013 (2020).

35 Botte, M., Ni, D., Schenck, S. et al. Cryo-EM structures of a LptDE transporter in complex with Pro-macrobodies offer insight into lipopolysaccharide translocation. Nature communications 13, 1826, doi:10.1038/s41467-022-29459-2 (2022).

36 Yang, Y., Chen, H., Corey, R. A. et al. LptM promotes oxidative maturation of the lipopolysaccharide translocon by substrate binding mimicry. Nature communications 14, 6368, doi:10.1038/s41467-023-42007-w (2023).

37 Sperandeo, P., Cescutti, R., Villa, R. et al. Characterization of lptA and lptB, two essential genes implicated in lipopolysaccharide transport to the outer membrane of Escherichia coli. Journal of bacteriology 189, 244–253, doi:10.1128/jb.01126-06 (2007).

38 Sueki, A., Stein, F., Savitski, M. M. et al. Systematic Localization of Escherichia coli Membrane Proteins. mSystems 5, doi:10.1128/mSystems.00808-19 (2020).

39 Jumper, J., Evans, R., Pritzel, A. et al. Highly accurate protein structure prediction with AlphaFold. Nature 596, 583–589, doi:10.1038/s41586-021-03819-2 (2021).

40 Bishop, R. E. The bacterial lipocalins. Biochimica et biophysica acta 1482, 73–83, doi:10.1016/s0167-4838(00)00138-2 (2000).

41 Gennaris, A., Nguyen, V. S., Thouvenel, L. et al. Optimal functioning of the Lpt bridge depends on a ternary complex between the lipocalin YedD and the LptDE translocon. Cell reports 44, 115446, doi:10.1016/j.celrep.2025.115446 (2025).

42 Schulz, G. E. The structure of bacterial outer membrane proteins. Biochimica et biophysica acta 1565, 308–317, doi:10.1016/s0005-2736(02)00577-1 (2002).

43 Fiorentino, F., Sauer, J. B., Qiu, X. et al. Dynamics of an LPS translocon induced by substrate and an antimicrobial peptide. Nature chemical biology 17, 187–195, doi:10.1038/s41589-020-00694-2 (2021).

44 Abraham, M., Schulz, Pall, Smith, Hessa, Lindahla. GROMACS: High performance molecular simulations through multi-level parallelism from laptops to supercomputers. SoftwareX, 10.1016/j.softx.2015.06.001 (2015).

45 Malojčić, G., Andres, D., Grabowicz, M. et al. LptE binds to and alters the physical state of LPS to catalyze its assembly at the cell surface. Proceedings of the National Academy of Sciences of the United States of America 111, 9467–9472, doi:10.1073/pnas.1402746111 (2014).

46 Sousa, M. C. New antibiotics target the outer membrane of bacteria. Nature 576, 389–390, doi:10.1038/d41586-019-03730-x (2019).

47 Robinson, J. A. Folded Synthetic Peptides and Other Molecules Targeting Outer Membrane Protein Complexes in Gram-Negative Bacteria. Frontiers in chemistry 7, 45, doi:10.3389/fchem.2019.00045 (2019).

48 Sperandeo, P., Martorana, A. M., Zaccaria, M. et al. Targeting the LPS export pathway for the development of novel therapeutics. Biochimica et biophysica acta. Molecular cell research 1870, 119406, doi:10.1016/j.bbamcr.2022.119406 (2023).

49 Lomize, M. A., Pogozheva, I. D., Joo, H. et al. OPM database and PPM web server: resources for positioning of proteins in membranes. Nucleic acids research 40, D370–376, doi:10.1093/nar/gkr703 (2012).

## SUPPLEMENTARY REFERENCES

1 Grenier, F., Matteau, D., Baby, V. et al. Complete Genome Sequence of Escherichia coli BW25113. Genome announcements 2, doi:10.1128/genomeA.01038-14 (2014).

2 Baba, T., Ara, T., Hasegawa, M. et al. Construction of Escherichia coli K-12 in-frame, single-gene knockout mutants: the Keio collection. Molecular systems biology 2, 2006.0008, doi:10.1038/msb4100050 (2006).

3 Yang, Y., Chen, H., Corey, R. A. et al. LptM promotes oxidative maturation of the lipopolysaccharide translocon by substrate binding mimicry. Nature communications 14, 6368, doi:10.1038/s41467-023-42007-w (2023).

4 Ranava, D., Yang, Y., Orenday-Tapia, L. et al. Lipoprotein DolP supports proper folding of BamA in the bacterial outer membrane promoting fitness upon envelope stress. Elife 10, doi:10.7554/eLife.67817 (2021).

5 Perez-Riverol, Y., Bai, J., Bandla, C. et al. The PRIDE database resources in 2022: a hub for mass spectrometry-based proteomics evidences. Nucleic acids research 50, D543–d552, doi:10.1093/nar/gkab1038 (2022).

6 Wilm, M., Shevchenko, A., Houthaeve, T. et al. Femtomole sequencing of proteins from polyacrylamide gels by nano-electrospray mass spectrometry. Nature 379, 466–469, doi:10.1038/379466a0 (1996).

7 Bouyssié, D., Dubois, M., Nasso, S. et al. mzDB: a file format using multiple indexing strategies for the efficient analysis of large LC-MS/MS and SWATH-MS data sets. Molecular & cellular proteomics : MCP 14, 771–781, doi:10.1074/mcp.O114.039115 (2015).

8 Bouyssié, D., Hesse, A. M., Mouton-Barbosa, E. et al. Proline: an efficient and user-friendly software suite for large-scale proteomics. Bioinformatics (Oxford, England) 36, 3148–3155, doi:10.1093/bioinformatics/btaa118 (2020).

9 Schwanhäusser, B., Busse, D., Li, N. et al. Global quantification of mammalian gene expression control. Nature 473, 337–342, doi:10.1038/nature10098 (2011).

10 Smits, A. H., Jansen, P. W., Poser, I. et al. Stoichiometry of chromatin-associated protein complexes revealed by label-free quantitative mass spectrometry-based proteomics. Nucleic acids research 41, e28, doi:10.1093/nar/gks941 (2013).

11 Mastronarde, D. N. Automated electron microscope tomography using robust prediction of specimen movements. Journal of structural biology 152, 36–51, doi:10.1016/j.jsb.2005.07.007 (2005).

12 Punjani, A., Rubinstein, J. L., Fleet, D. J. et al. cryoSPARC: algorithms for rapid unsupervised cryo-EM structure determination. Nature methods 14, 290–296, doi:10.1038/nmeth.4169 (2017).

13 Jumper, J., Evans, R., Pritzel, A. et al. Highly accurate protein structure prediction with AlphaFold. Nature 596, 583–589, doi:10.1038/s41586-021-03819-2 (2021).

14 Croll, T. I. ISOLDE: a physically realistic environment for model building into low-resolution electron-density maps. Acta crystallographica. Section D, Structural biology 74, 519–530, doi:10.1107/s2059798318002425 (2018).

15 Emsley, P., Lohkamp, B., Scott, W. G. et al. Features and development of Coot. Acta crystallographica. Section D, Biological crystallography 66, 486–501, doi:10.1107/s0907444910007493 (2010).

16 Sanchez-Garcia, R., Gomez-Blanco, J., Cuervo, A. et al. DeepEMhancer: a deep learning solution for cryo-EM volume post-processing. Communications biology 4, 874, doi:10.1038/s42003-021-02399-1 (2021).

17 Liebschner, D., Afonine, P. V., Baker, M. L. et al. Macromolecular structure determination using X-rays, neutrons and electrons: recent developments in Phenix. Acta crystallographica. Section D, Structural biology 75, 861–877, doi:10.1107/s2059798319011471 (2019).

18 Williams, C. J., Headd, J. J., Moriarty, N. W. et al. MolProbity: More and better reference data for improved all-atom structure validation. Protein science : a publication of the Protein Society 27, 293–315, doi:10.1002/pro.3330 (2018).

19 Abraham, M., Schulz, Pall, Smith, Hessa, Lindahla. GROMACS: High performance molecular simulations through multi-level parallelism from laptops to supercomputers. SoftwareX, 10.1016/j.softx.2015.06.001 (2015).

20 Stansfeld, P. J., Goose, J. E., Caffrey, M. et al. MemProtMD: Automated Insertion of Membrane Protein Structures into Explicit Lipid Membranes. Structure (London, England : 1993) 23, 1350–1361, doi:10.1016/j.str.2015.05.006 (2015).

21 Souza, P. C. T., Alessandri, R., Barnoud, J. et al. Martini 3: a general purpose force field for coarse-grained molecular dynamics. Nature methods 18, 382–388, doi:10.1038/s41592-021-01098-3 (2021).

22 de Jong, D. H., Singh, G., Bennett, W. F. et al. Improved Parameters for the Martini Coarse-Grained Protein Force Field. Journal of chemical theory and computation 9, 687–697, doi:10.1021/ct300646g (2013).

23 Huang, J., Rauscher, S., Nawrocki, G. et al. CHARMM36m: an improved force field for folded and intrinsically disordered proteins. Nature methods 14, 71–73, doi:10.1038/nmeth.4067 (2017).

24 Vickery, O. N. & Stansfeld, P. J. CG2AT2: an Enhanced Fragment-Based Approach for Serial Multi-scale Molecular Dynamics Simulations. Journal of chemical theory and computation 17, 6472–6482, doi:10.1021/acs.jctc.1c00295 (2021).

25 Bernetti, M. & Bussi, G. Pressure control using stochastic cell rescaling. The Journal of chemical physics 153, 114107, doi:10.1063/5.0020514 (2020).

26 Bussi, G., Donadio, D. & Parrinello, M. Canonical sampling through velocity rescaling. The Journal of chemical physics 126, 014101, doi:10.1063/1.2408420 (2007).

27 Michaud-Agrawal, N., Denning, E. J., Woolf, T. B. et al. MDAnalysis: a toolkit for the analysis of molecular dynamics simulations. Journal of computational chemistry 32, 2319–2327, doi:10.1002/jcc.21787 (2011).

28 Gowers, R. J., Linke, M., Barnoud, J. et al. MDAnalysis: A Python Package for the Rapid Analysis of Molecular Dynamics Simulations. Proceedings of the Python in Science Conference, doi:10.25080/Majora-629e541a-00e (2019/09/11).

29 Parks, D. H., Chuvochina, M., Rinke, C. et al. GTDB: an ongoing census of bacterial and archaeal diversity through a phylogenetically consistent, rank normalized and complete genome-based taxonomy. Nucleic acids research 50, D785–d794, doi:10.1093/nar/gkab776 (2022).

30 Seemann, T. Prokka: rapid prokaryotic genome annotation. Bioinformatics (Oxford, England) 30, 2068–2069, doi:10.1093/bioinformatics/btu153 (2014).

31 Emms, D. M. & Kelly, S. OrthoFinder: phylogenetic orthology inference for comparative genomics. Genome biology 20, 238, doi:10.1186/s13059-019-1832-y (2019).

32 Katoh, K. & Standley, D. M. MAFFT multiple sequence alignment software version 7: improvements in performance and usability. Molecular biology and evolution 30, 772–780, doi:10.1093/molbev/mst010 (2013).

33 Minh, B. Q., Schmidt, H. A., Chernomor, O. et al. IQ-TREE 2: New Models and Efficient Methods for Phylogenetic Inference in the Genomic Era. Molecular biology and evolution 37, 1530–1534, doi:10.1093/molbev/msaa015 (2020).

34 Letunic, I. & Bork, P. Interactive Tree Of Life (iTOL) v5: an online tool for phylogenetic tree display and annotation. Nucleic acids research 49, W293–w296, doi:10.1093/nar/gkab301 (2021).

35 Adeolu, M., Alnajar, S., Naushad, S. et al. Genome-based phylogeny and taxonomy of the ‘Enterobacteriales’: proposal for Enterobacterales ord. nov. divided into the families Enterobacteriaceae, Erwiniaceae fam. nov., Pectobacteriaceae fam. nov., Yersiniaceae fam. nov., Hafniaceae fam. nov., Morganellaceae fam. nov., and Budviciaceae fam. nov. International journal of systematic and evolutionary microbiology 66, 5575–5599, doi:10.1099/ijsem.0.001485 (2016).

36 Parte, A. C., Sardà Carbasse, J., Meier-Kolthoff, J. P. et al. List of Prokaryotic names with Standing in Nomenclature (LPSN) moves to the DSMZ. International journal of systematic and evolutionary microbiology 70, 5607–5612, doi:10.1099/ijsem.0.004332 (2020).

37 Finn, R. D., Clements, J. & Eddy, S. R. HMMER web server: interactive sequence similarity searching. Nucleic acids research 39, W29–37, doi:10.1093/nar/gkr367 (2011).

38 Capella-Gutiérrez, S., Silla-Martínez, J. M. & Gabaldón, T. trimAl: a tool for automated alignment trimming in large-scale phylogenetic analyses. Bioinformatics (Oxford, England) 25, 1972–1973, doi:10.1093/bioinformatics/btp348 (2009).

39 Kalyaanamoorthy, S., Minh, B. Q., Wong, T. K. F. et al. ModelFinder: fast model selection for accurate phylogenetic estimates. Nature methods 14, 587–589, doi:10.1038/nmeth.4285 (2017).

40 Chin, J. W., Martin, A. B., King, D. S. et al. Addition of a photocrosslinking amino acid to the genetic code of Escherichiacoli. Proceedings of the National Academy of Sciences of the United States of America 99, 11020–11024, doi:10.1073/pnas.172226299 (2002).

41 Chaveroche, M. K., Ghigo, J. M. & d’Enfert, C. A rapid method for efficient gene replacement in the filamentous fungus Aspergillus nidulans. Nucleic acids research 28, E97, doi:10.1093/nar/28.22.e97 (2000).

